# Unraveling dendritic-cell diversity in blood of pigs: welcome tDC and DC3

**DOI:** 10.1101/2025.03.21.644143

**Authors:** Ambre Baillou, Gaël Auray, Francisco Brito, Marius Botos, Alizée Huber, Artur Summerfield, Stephanie C. Talker

## Abstract

Dendritic cells (DC) are professional antigen presenting cells playing a major role in orchestrating adaptative immune responses. To adapt to various immune challenges, such as different classes of pathogens, specialized subsets of DC have evolved across species. To date, DC are classified as conventional DC (cDC1, cDC2) and plasmacytoid DC (pDC), with the more recent addition of DC3 and transitional DC (tDC) that were discovered in human and mouse thanks to high-dimensional phenotyping and single-cell sequencing technologies.

Here, by combining flow cytometry and RNA-seq on the bulk- and single-cell level, we identified the porcine equivalent of tDC in blood as CD14^-^CADM1^-^CD172a^+^CD4-cells expressing both Flt3 and CD123 (IL-3RA). This new subset forms a well-defined cluster when mapped onto scRNA-seq data of enriched DC and shares transcriptomic features and abundance with porcine blood cDC2 and pDC. Moreover, we describe putative porcine DC3 as transcriptionally overlapping cells in-between cDC2 and monocytes. With the core functions of tDC and DC3 remaining to be elucidated, our datasets provide a valuable resource for cross-species research on DC heterogeneity in various lymphoid and non-lymphoid tissues.

## INTRODUCTION

Dendritic cells (DC) are best known as instructors of T-cell immunity through antigen presentation and co-stimulation. Their response enables the system to adapt to various challenges and simultaneously ensures tolerance to harmless antigens. To fulfill these diverse roles, the DC system comprises phenotypically and functionally distinct cell subsets that have been extensively studied in humans (1,2), mice (3,4) and pigs (5,6) among other species (7–11). Traditionally, DC have been broadly divided into two lineages: plasmacytoid DC (pDC), primarily known as IFN type 1 producers in response to viral infection, and conventional DC (cDC) which are highly efficient in stimulating T-cell responses. Conventional DC were further divided into type 1 cDC (cDC1) and type 2 cDC (cDC2), with cDC1 appearing specialized in the induction of Th1- and cytotoxic T-cell responses, and cDC2 preferentially promoting Th2/Th17 responses (4). Over the last decade, high-dimensional and high-throughput approaches, such as single-cell RNA sequencing (scRNA-seq) (1), have revealed an astonishing heterogeneity and plasticity of cDC2 (1,12,13), with cDC2 subsets likely arising from distinct ontogenetic lineages (14–16) and being shaped by signals in their microenvironment (17). Moreover, highly pro-inflammatory cDC2 have been identified as a separate DC lineage, namely type 3 DC (DC3), overlapping with monocytes both phenotypically and transcriptionally and putting traditional monocyte markers like CD14 into question (13,15,18,19).

Furthermore, the phenotypic definition of pDC was challenged, when putative pre-DC were discovered with scRNA-seq in humans and mice and shown to contaminate traditional pDC gates(20–22). Indeed, these putative pre-DC, shown to derive from pro-pDC and now classified as transitional DC (tDC), appear to be competent antigen presenters that have likely biased several *in vitro* assays involving pDC, as discussed here: (20,21,23,24). A murine coronavirus infection model has suggested the involvement of tDC in viral responses, with the intriguing hypothesis that tDC are in a delicate balance with antiviral pDC and enhance pro-inflammatory responses by IL1-β production (25). Notably, cDC2-like cells were shown to differentiate from tDC, and may be termed tDC2 as suggested by Sulczewski *et al.* (25). These tDC2 very much resemble ESAM^+^ cDC2 and CD5^+^ cDC2 in mouse and human, respectively, and were shown to replenish the DC2 pool in mouse models with impaired pre-DC2 development (25–27). This further complicates the picture of DC2 heterogeneity, now encompassing pre-cDC-derived cDC2 subsets (16), pro-DC3-derived DC3 (monocyte-like) (19), and pro-pDC-derived tDC2 (pDC-like) (25).

We have previously described phenotype and bulk transcriptome of porcine blood cDC1, cDC2 and pDC, with key gene expression confirming a gating strategy based on CD14, CD172a, CADM1 and CD4 (5). Accordingly, porcine cDC1 can be identified as CD14-CD172a^low^CADM1^+^CD4^-^, cDC2 as CD14^-^CD172a^+^CADM1^+^CD4^-^, and pDC as CD14-CD172a^+^CADM1^-^CD4^+^. Notably, using this marker combination, one additional subset was apparent, expressing CD172a, but lacking expression of all other markers (CD14^-^CADM1-CD172a^+^CD4^-^). By performing scRNA-seq on Flt3^+^ DC enriched from porcine blood, we now confirm the existence of this novel subset and identify it as porcine tDC. Moreover, we describe putative DC3 as cells co-expressing both cDC2 markers (*FLT3*, *FCER1A*, *CD1.1*) and monocyte markers (*CSF1R*, *CD14*, *CD163*, *C5AR1*).

## MATERIAL AND METHODS

### Ethics statement

Blood sampling of pigs was performed in compliance with the Swiss animal protection law (TSchG SR 455; TSchV SR 455.1; TVV SR 455.163). The procedures were reviewed by the committee on animal experiments of the canton of Bern, Switzerland, and approved by the cantonal veterinary authority (Amt für Landwirtschaft und Natur LANAT, Veterinärdienst VeD, Bern, Switzerland) under the licence numbers BE88/14 and BE127/2020.

### Animals and isolation of PBMC

Blood was obtained from Swiss Large White pigs (**Table 1**) kept under specific-pathogen-free (SPF) conditions (28) at the animal facility of the IVI (Mittelhäusern, Switzerland) by puncturing the jugular vein. As anti-coagulant, citrate-based Alsever’s solution (1.55 mM C_6_H_12_O_6_; 408 mM Na_3_C_6_H_5_O_7_·2H_2_O; 1.078 mM NaCl; 43 mM C_6_H_8_O_7_, pH 6.2) was used.

**Table 1.** Animals used in each experimentation.

| Experiment | Number of animals<br>(Swiss Large White pigs) | Age | Male/Female |
| --- | --- | --- | --- |
| **bulk RNA-seq of sorted cDC1, cDC2, pDC, and mon** |  |  |  |
| (From Auray *et al.*, 2016) | n = 3 | 3-12 months | F |
| **bulk RNA-seq of sorted pptDC** |  |  |  |
|  | n = 4 | 12-24 months | F |
| **scRNA-seq of enriched DC** |  |  |  |
|  | n = 3 | 16.5 months | F |

For peripheral blood mononuclear cell (PBMC) isolation, blood was centrifuged at 1 000 x g for 20 min (room temperature; RT), the buffy coat was collected, diluted in PBS/EDTA (PBS; 1 mM EDTA) to a 1:1 ratio (RT) and layered onto Ficoll-paque (1.077 g/L, GE Healthcare) in Leucosep tubes (Greiner BioOne) for centrifugation at 800 x g for 25 min (RT). PBMC were collected and washed first once with cold PBS/EDTA at 350 x g for 10 min (+4°C, Ficoll-paque removal) and then once reducing the speed to 250 x g (platelet removal). Finally, red blood cells were removed from the PBMC by incubation with cold lysis buffer (10 mM NaHCO_3_; 1 mM EDTA; 0.15 M NH_4_Cl, pH 7.25) for 10 min on ice, followed by two washes with cold PBS/EDTA at 250 x g for 10 min (+4°C).

### Phenotyping of putative porcine tDC in blood by flow cytometry

The flow cytometry gating strategy used to identify mononuclear phagocyte (MP) subsets in pig blood was previously described by our laboratory (5), defining cDC1 as CD14-CD172a^low^CADM1^+^ cells, cDC2 as CD14^-^CD172a^+^CADM1^+^ cells, pDC as CD14-CD172a^+^CADM1^-^CD4^+^ cells and monocytes as CD14^+^ cells. The same staining panel was used to gate on the newly identified DC subset as CD14^-^CD172a^+^CADM1^-^CD4-cells in the present study. Briefly, a four-step four-color staining of PBMC was performed, with two washes (400 x g, 4 min, 4°C) with Cell Wash (BD Biosciences) in-between incubations. Antibodies and porcine recombinant proteins used are listed in **Table 2**. Briefly, cells were first stained with the primary antibodies anti-CD172a (clone 77-22-15A) and anti-SynCAM (TSLC1/CADM1, clone 3E1), followed by a second incubation with the corresponding secondary antibodies anti-mouse IgG2b AF647 and anti-chicken IgY biotin. A blocking step was then performed with ChromPure mouse IgG (Jackson Immunoresearch), and cells were finally incubated with the directly conjugated antibodies anti-CD14-FITC (clone MIL2) and anti-CD4-PerCP-Cy5.5 (clone 74-12-4), and with V500-conjugated streptavidin. Based on this staining, the phenotype of the new DC subset of interest was further characterized by analyzing the expression of additional cell surface markers, alongside corresponding FMO (Fluorescence minus one) controls. All the flow cytometry acquisitions were performed on a FACS Canto II (BD Bioscience) equipped with three lasers (405, 488, and 633 nm), using the DIVA software and were further analyzed with the Flowjo software (TreeStar, version 10.10.0).

**Table 2:** List of antibodies and porcine recombinant proteins for phenotyping and FACS.

| Experiment | Antigen or receptor* | Clone/Source of mAb | Detection/Source |
| --- | --- | --- | --- |
| **Phenotyping<sup>1</sup>/**<br>**FACS for bulk RNA-seq<sup>2</sup>** |  |  |  |
| Core | CD172a | 77-22-15A/VetMedUni Vienna, Austria | Anti-mouse IgG2b:A647/Molecular Probes |
|  | CADM1** | 3E1/MBL | Anti-chicken IgY:biotin/Jackson ImmunoResearch<br>+ V500-coupled streptavidin/BD Horizon<sup>1</sup> or<br>+ A750-APC-coupled streptavidin/Thermo Fisher<sup>2</sup> |
|  | CD14 | MIL2:FITC/AbD Serotec |  |
|  | CD4 | 74-12-4:PerCP-Cy5.5/BD Pharmingen |  |
| Phenotypic markers | wC11R1/CD11b | MIL4/Serotec | Anti-mouse IgG1:RPE/SouthernBiotech |
|  | CD1.1 | 76-7-4/VetMedUni Vienna, Austria | Anti-mouse IgG2a:RPE/SouthernBiotech |
|  | CD115/CSF1R | ROS8G11-1/Roslin Institute, University of Edinburgh, UK | Anti-mouse IgG2a:RPE/SouthernBiotech |
|  | CD163 | 2A10-11/INIA-CSIC, Madrid, Spain | Anti-mouse IgG1:RPE/SouthernBiotech |
|  | CD205 | ZH9F7/CIAD, Hermosillo, Mexico | Anti-mouse IgG1:RPE/SouthernBiotech |
|  | CD303 | 102G7/Dendritics, Lyon France | Anti-mouse IgG1:RPE/SouthernBiotech |
|  | MHC-II/SLA-DQ | TH16B/VMRD | Anti-mouse IgG2a:RPE/SouthernBiotech |
|  | CD16 | G7:RPE/AdB Serotec |  |
|  | CD135/Flt3* | His-tagged porcine recombinant protein Flt3L/In house | Anti-His:PE/Miltenyi Biotec |
|  | CD123/IL3RA* | His-tagged porcine recombinant protein IL-3/In house | Anti-His:RPE/Miltenyi Biotec |
| T-cell depletion | CD3 | PTT3-FyH2/University of Bristol, UK | Anti-mouse IgG:magnetic beads/Miltenyi Biotec |
| **DC enrichment for scRNA-seq** |  |  |  |
| Core | CD172a | 77-22-15A/VetMedUni Vienna, Austria | Anti-mouse IgG2b:A647/Molecular Probes |
|  | CD135/Flt3* | His-tagged porcine recombinant protein Flt3L/In house | Anti-His:PE/Miltenyi Biotec |
| T-cell depletion | CD3 | 8E6-8C8/Kingfisher Biotech | Anti-mouse IgG2a:biotin<br>+ Magnetic beads:streptavidin/Miltenyi Biotec |

\*\* Anti-mouse CADM1 with pig cross reactivity

### Sorting and bulk RNA sequencing of putative porcine blood tDC

The newly identified DC subset was sorted from the blood of four pigs (12 to 24-month-old) for bulk RNA-seq analysis. First, a T-cell depletion of PBMC was performed using magnetic activated cell sorting with an anti-CD3 antibody (clone 8E6-8C8), anti-mouse IgG MicroBeads and LD columns (MACS MicroBead Technology, Miltenyi Biotec). The same four-step four-color staining as described above was performed with the CD3-negative fraction, but V500-conjugated streptavidin was replaced by A750-APC streptavidin, and the DC subset of interest was sorted using fluorescence activated cell sorting (FACS; FACSAria III; BD Biosciences). Finally, cells were resuspended in TRIzol (Life Technologies) and stored at −80°C until later RNA extraction with the Nucleospin RNA kit (Macherey Nagel) as previously described (5). RNA quantification and quality assessment was performed with an Agilent 2100 Bioanalyzer (Agilent Technologies) and a Qubit 2.0 Fluorometer (Life Technologies). High-quality RNA (approximately 500 ng; RNA integrity number (RIN) > 8) was used to prepare non-directional paired-end mRNA libraries with the TruSeq Sample Preparation Kit (v2, Illumina). The libraries were sequenced on the Illumina HiSeq2500 platform using 2 x 100 bp paired-end sequencing cycles, yielding between 26.5 and 30.1 million read pairs per sample. The Illumina BCL output files with base calls and qualities were converted into FASTQ file format and demultiplexed with the CASAVA software (v1.8.2). Raw bulk RNA-seq data for the cDC1, cDC2, pDC and monocytes (n = 3 pigs) were available from our previous work (5).

For analysis of bulk RNA-seq data, the following bioinformatics tools were used with their default parameters, unless specified otherwise. The quality of reads was assessed with fastQC v0.11.9 (https://www.bioinformatics.babraham.ac.uk/projects/fastqc/) and both low quality bases (Phred score < 30) and Illumina TruSeq2 adapters were trimmed with Trimmomatic v0.39 (29). The reads were then mapped to the pig reference genome (assembly Sscrofa 11.1) using STAR v2.7.10a (30). The duplicate reads were identified and removed using MarkDuplicates from the Picard command-line tools v2.25.1 (https://broadinstitute.github.io/picard/). The featureCounts program included in the SourceForge Subread package v2.0.3 (31) was used to count the number of reads overlapping with each gene identified in the Ensembl pig annotation release 11.1.111. In summary, (i) a minimum of 93.7% of reads were mapped to the genome among all samples, yielding between 29.6 and 57.2 million reads aligned per sample, (ii) between 14.2 and 35.4% of reads were identified as duplicates and (iii) 58.5-68.9% of reads were assigned to a gene, corresponding to a range of 16.3-33.0 million mapped reads.

The differential gene expression analyses were performed using the Bioconductor package DESeq2 v1.42.1 (32) in R v4.3.3 (33). Only genes with |log2FC| > 1 and adjusted p-value < 0.05 were selected as differentially expressed genes (DEGs). We performed pairwise comparisons of cell subsets as well as the comparison of each subset versus all others to define the MP subset-specific transcriptomic signatures (results are available as **Supp. Data 1**). Principal component analysis (PCA) was performed with normalized and vst-transformed counts of the 500 most variable genes across samples. Sample-sample correlation analysis was based on normalized gene expression data for each sample (*counts()*) using the Spearman correlation coefficient with hierarchical clustering based on Spearman distances.

### Enrichment of DC by fluorescence-activated cell sorting (FACS)

To enrich DC for scRNA-seq analysis, a four-step protocol combining cell staining and T-cell depletion was performed on freshly isolated PBMC from three pigs in parallel. In-between incubation steps, cells were washed twice (400 x g, 4 min, 4°C) with BD Cell Wash (BD Biosciences). Antibodies and porcine recombinant proteins used are listed in **Table 2**. Briefly, 5 x 10^8^ PBMC were first stained with a His-tagged porcine recombinant protein Flt3L and the primary antibodies anti-CD172a (clone 77-22-15A) and anti-CD3 (clone 8E6-8C8), followed by a second incubation step with the corresponding secondary antibodies anti-His-PE, anti-mouse IgG2b AF647 and anti-mouse IgG2a biotin. Next, following incubation with Streptavidin MicroBeads (Miltenyi Biotec), CD3^+^ cells were depleted using LD columns (Miltenyi Biotec). Finally, total DC from the three pigs identified as FLT3^+^CD172a^-/+^ cells were sorted in parallel using one FACS Aria II and two FACS Aria III (all BD Bioscience) at the flow cytometry and cell sorting core facility at the University of Bern.

### Single-cell RNA-seq (10x Genomics)

For scRNA-seq, DC isolated from the blood of three pigs (16.5-month-old) were analyzed with 10x Genomics. Gel beads-in-emulsion (GEM) generation and barcoding, reverse transcription, cDNA amplification and 3’ gene expression library generation steps were all performed according to the Chromium Next GEM Single Cell 3ʹ Reagent Kits v3.1 (Dual Index) User Guide (10x Genomics CG000315, Rev E) with all stipulated 10x Genomics reagents. Generally, 9-11 µL of each cell suspension (1 500-1 900 cells/µL) and 32-35 µL of nuclease-free water were used for a targeted cell recovery of 10 000 cells. GEM generation was followed by a GEM-reverse transcription incubation, a clean-up step and 11 cycles of cDNA amplification. The resulting cDNA was evaluated for quantity and quality using a Thermo Fisher Scientific Qubit 4.0 fluorometer with the Qubit dsDNA HS Assay Kit (Thermo Fisher Scientific, Q32851) and an Advanced Analytical Fragment Analyzer System using a Fragment Analyzer NGS Fragment Kit (Agilent, DNF-473), respectively. Thereafter, 3ʹ sc gene expression libraries were constructed using a sample index PCR step of 14 cycles. The generated cDNA libraries were tested for quantity and quality using fluorometry and capillary electrophoresis as described above. The cDNA libraries were pooled and sequenced with a loading concentration of 300 pM, asymmetric paired-end and dual indexed, on two shared Illumina NovaSeq 6000 sequencer using a NovaSeq 6000 S4 Reagent Kits v1.5 (200 cycles; Illumina, 20028313). The read set-up was as follows: read 1: 29 cycles, i7 index: 10 cycles, i5: 10 cycles and read 2: 91 cycles. The quality of the sequencing runs was assessed using Illumina Sequencing Analysis Viewer (v2.4.7, Illumina) and all base call files were demultiplexed and converted into FASTQ files using bcl2fastq conversion software (v2.20, Illumina). The mean reads per cell and number of cells obtained per sample ranged from 33 896 to 51 718 reads, and from 11 5311 to 14 221 cells, respectively All steps were performed at the Next Generation Sequencing Platform, University of Bern.

### Analysis of porcine scRNA-seq data

#### Read alignment, quality control and filtering

The scRNA-seq FASTQ files were processed using Cell Ranger v7.1.0 (10x Genomics) (34) and reads were aligned to the pig reference genome (assembly Sscrofa 11.1). Bam files and filtered expression matrices were generated using the “cellranger_count” pipeline with default parameters, unless specified otherwise. Expression matrices were further analyzed in R v4.3.3 (33) using mainly Seurat v5.1.0 (35) and other R packages (list available in the GitHub page, see **“Code Availability”** section). Quality-based scRNA-seq data filtering was performed by excluding low-quality cells and dead cells (< 500 genes and > 10% of transcripts mapping to mitochondrial genes), non-expressed genes (genes expressed in < 5 cells across all samples) and cells identified with high probability as doublet by the scDblFinder package v1.16.0 (36) (doublet score threshold automatically determined). Percentages of mitochondrial and ribosomal protein gene expression in cells were calculated based on *ND1*, *ND2*, *COX1*, *COX2*, *ATP8*, *ATP6*, *COX3*, *ND3*, *ND4L*, *ND4*, *ND5*, *ND6*, *CYTB* genes and 61 *RPS-* and *RPL-* genes, respectively.

#### Normalization, dimensionality reduction, data integration and clustering

The three scRNA-seq samples loaded in a Seurat object (1 layer/sample) were independently processed for sctransform-based normalization, including steps of data scaling and highly variable gene identification, and for linear dimensionality reduction using PCA. For further downstream analysis, the optimal number of 50 principal components (PCs) was identified by the elbowplot method. Cells were then scored for cell cycle phases based on their expression of S and G2M phase-associated genes listed in Seurat. Next, data integration of the three datasets was performed with Harmony v1.2.0 (37) on PCA cell embeddings and selecting the sample origin (batch effect correction) and the cell cycle phase as covariates. The resulting Harmony reduction was selected for identifying nearest neighbors, clustering the cells with the Leiden algorithm (method = “*igraph*”, clustering resolution = 0.6), and performing non-linear dimensional reduction using UMAP for cluster visualization.

#### Differential gene expression analysis

Counts in the RNA assay were log-normalized and scaled, and layers were joined. The DEGs in each cluster were identified with the *FindAllMarkers()* function and pairwise comparisons between selected clusters were performed with the *FindMarkers()* function. Only genes expressed in at least 20% of the cells in one of the clusters being compared, with |avg_log2FC| > 1 and adjusted p-value < 0.05 were selected as DEGs. In addition, DEGs from pairwise comparisons were filtered according to an expression in at least 20% of the cells in one of the clusters being compared and 80% of the cells in the other one(s).

#### Cluster correlation analysis

Cluster-cluster correlation values were calculated based on averaged log-normalized gene expression data for each cluster using the Spearman correlation coefficient.

#### Gene set enrichment analysis

Gene set enrichment analyses (GSEA) were performed with the AUCell package v1.24.0 (38) as previously described by Herrera-Uribe *et al*. (39). Briefly, the expression of the specific enriched gene set in each sorted porcine blood MP subset analyzed by bulk RNA-seq (as described in preceding methods) was evaluated within cells of the scRNA-seq dataset, as follows: Ranking of gene expression from raw gene counts and calculation of area under the curve (AUC) scores from the top 5, 10, 15, 25, 50 and 100% of expressed genes in a cell and the gene sets. AUC scores are proportional to the percentage of genes from a gene set found in the top expressed genes for a cell defined at different levels. Next, AUC scores and UMAP coordinates of each cell were overlayed for UMAP visualization, with manual determination of a threshold value for each gene set based on AUC score distributions. Heatmap representation was based on averaged scaled AUC scores calculated for each cluster, following scaling of individual cell AUC scores relative to other cells within a single gene set comparison (rows) but not between gene sets (columns).

For species comparison, GSEA were performed with DC subset gene signatures from three sources: (i) a published scRNA-seq study (SMARTSeq2) of human blood DC (20), (ii) a published bulk and scRNA-seq study of murine spleen DC (10x Genomics) (25), and (iii) results from re-analysis of a recently published scRNA-seq dataset (10x Genomics) of human blood DC (40). The pig orthologs of human and mouse genes were identified with BioMart (Ensembl) and selected according to the highest percentage of identity to the target pig gene. Human and mouse genes without pig orthologs were removed. The resulting pig-converted-human and murine gene signatures are provided as **Supplementary Data 2**. GSEA were performed as described above, calculating the AUC scores from the top 25% of expressed genes in a cell and the gene sets and represented as heatmaps.

#### Machine learning-based cell scoring

The classification score for the different cell clusters was created with the scikit-learn python module (41) as previously described by Mayère *et al.* (42). An ElasticNet model with one versus all approach was trained using the *ElasticNet()* function (alpha = 0.05, tol = 0.01) on a random subsample of 450 cells per cluster in order to avoid gene weighting bias due to overrepresentation of some clusters. The ElasticNet approach uses a linear regression with combined L1 (Lasso) and L2 (Ridge) priors as regularizer, allowing a robust selection of relevant genes defining the cells of interest (43,44).

#### Cluster subsetting

Subsetting was performed using the Seurat’s *subset()* function. Non-expressed genes in the new datasets were removed with *DietSeurat()* and data were split according to the sample of origin using the *split()* function (1 layer/sample). Data were then re-processed for normalization, dimensional reduction, data integration and clustering (method = “*igraph*”) as described above. Counts in the RNA assay were log-normalized and scaled, and layers were joined.

#### Trajectory inference analysis

The trajectory inference analysis of the subsetted dataset was performed with the Scorpius package v1.0.9 (45). Cells were ordered according to the inferred linear trajectory using the *infer_trajectory()* function and the importance of a gene and its expression with respect to the modelled dynamic process was assessed with the *gene_importances()* function. Next, the top 50 important genes were assigned into modules according to their expression patterns across the inferred trajectory with the *extract_modules()* function, using the normalized expression values scaled from 0 to 1 with the *scale_quantile()* function.

### Analysis of published scRNA-seq data (human DC)

We analyzed the scRNA-seq dataset of human blood DC recently generated by Lubin *et al.* (40) (approximately 3 000 cells). DC were sorted by flow cytometry and subjected to 10x Genomics scRNA-seq. Processed data from the cellranger pipeline (barcode, feature and matrix files), available under the sample number GSM8499782 in the National Center for Biotechnology Information Gene Expression Omnibus database, were analyzed with the Seurat pipeline as described above for the porcine data (quality-based data filtering, sctransform-based normalization and linear dimensional reduction using PCA). The PCA reduction was selected for identifying nearest neighbors, clustering the cells with the Leiden algorithm (method = “*igraph*”, clustering resolution = 0.8), and performing non-linear dimensional reduction using UMAP for cluster visualization. Next, the differential gene expression analysis was performed as for the porcine data, using *FindAllMarkers()* to identify the DEGs in each cluster.

### Identification and replacement of gene identifiers

Pig gene Ensembl stable identifiers (IDs) without available gene name/symbol in the pig genome annotation file were replaced in text and figures by NCBI gene (formerly Entrezgene) accession or UniProtKB Gene Name symbol if available in the corresponding databases using the BioMart data mining tool from Ensembl (https://www.ensembl.org/biomart/martview). A list of replaced Ensembl IDs is included in **Supplementary Data 1**. The human gene names *HLA-DRA* and *HLA-DOB* found in the pig genome annotation were replaced by *SLA-DRA* and *SLA-DOB* respectively, the gene names of their porcine orthologs. While *IL3RA* is not currently annotated in the Ensembl pig genome, it is available in the NCBI reference. Thus, the porcine genomic sequence for the gene encoding IL3RA (ENSSSCG00000055271) was identified by aligning the *IL3RA* gene sequence from NCBI (gene identifier: 102166116) to the Ensembl pig genome (Sscrofa release 11.1.111) using the Ensembl BLAT (100% sequence identity). The sequence of the *IL3RA* transcript ENSSSCT00000092699, product of the ENSSSCG00000055271 gene (*IL3RA*), was utilized to visualize the read distribution across its corresponding genomic location (AEMK02000569.1: 775,837-784,610) for each sorted MP subset, using Integrative Genomics Viewer (IGV) software.

### Preparation of Figures

Figures were prepared using FlowJoTM v10.10.0 (BD Life Sciences) (46), R v4.3.3 (33), Rstudio v2024.04.1 (47), Inkscape v1.3.2 (https://www.inkscape.org), Integrative Genomics Viewer (IGV) v2.17.4 (48) softwares. FACS scheme was created using Servier Medical Art, by Servier (http://smart.servier.com).

Bulk RNA-seq data was represented as PCA and heatmaps using the ggplot2 v3.5.1 (49) and ComplexHeatmap v2.18.0 (50) R packages, respectively. Heatmaps were prepared following log10 transformation of normalized counts. Prior to log10 transformation, a pseudocount of 1 was added to the values to avoid zeros.

Visualization of scRNA-seq data was based on feature plots, dot plots, violin plots, bar plots, scatter plots and heatmaps using the Seurat v5.1.0 (35), scCustomize v2.1.2 (51), ggplot2 v3.5.1 (49) and ComplexHeatmap v2.18.0 (50) R packages. Heatmaps were generated with scaled and centered data (Seurat *ScaleData() function*). For improved contrast in feature plots, feature-specific contrast levels were calculated based on quantiles (q10, q90) of non-zero expression. Code is available from the visualization vignette of the Seurat package (https://www.satijalab.org/seurat/).

The cell classification scoring based on a machine learning model was visualized by scatter plots using the scikit-learn python module (41).

The trajectory inference analysis was represented as UMAP plot and heatmap using the Scorpius R package v1.0.9 (45).

## RESULTS

### Phenotype of putative porcine tDC (pptDC)

In a previous study, we characterized mononuclear phagocyte (MP) subsets in porcine blood by flow cytometry, identifying cDC1 as CD14^-^CD172a^low^CADM1^+^, cDC2 as CD14-CD172a^+^CADM1^+^, pDC as CD14^-^CD172a^+^CADM1^-^CD4^+^ and monocytes as CD14^+^ (5). Here, applying the same staining protocol and gating strategy, we focused our analysis on the previously undescribed MP subset of CD14^-^CD172a^+^ cells lacking both CADM1 and CD4 expression and being as frequent as pDC in blood of pigs (**Fig. 1A**). Expression of the conserved DC marker Flt3 (CD135), together with the almost complete lack of monocyte markers (CD115/CSF1R and CD163), supported its identification as a bona fide DC subset (**Fig. 1B**). Notably, it shared phenotypic markers with both cDC (CD11b/wCD11R1 (52) and CD205, absence of CD303) and pDC (IL3-RA/CD123, absence of CD1). Moreover, this subset expressed CD80/86 and a high level of MHC-II, thus suggesting its involvement in antigen presentation and T-cell co-stimulation. Taken together, this new DC subset displayed a phenotypic profile overlapping with both cDC and pDC phenotypes, and it shared prominent expression of IL3-RA with pDC, leading us to further address this subset as putative porcine tDC (pptDC).

**Figure 1:**
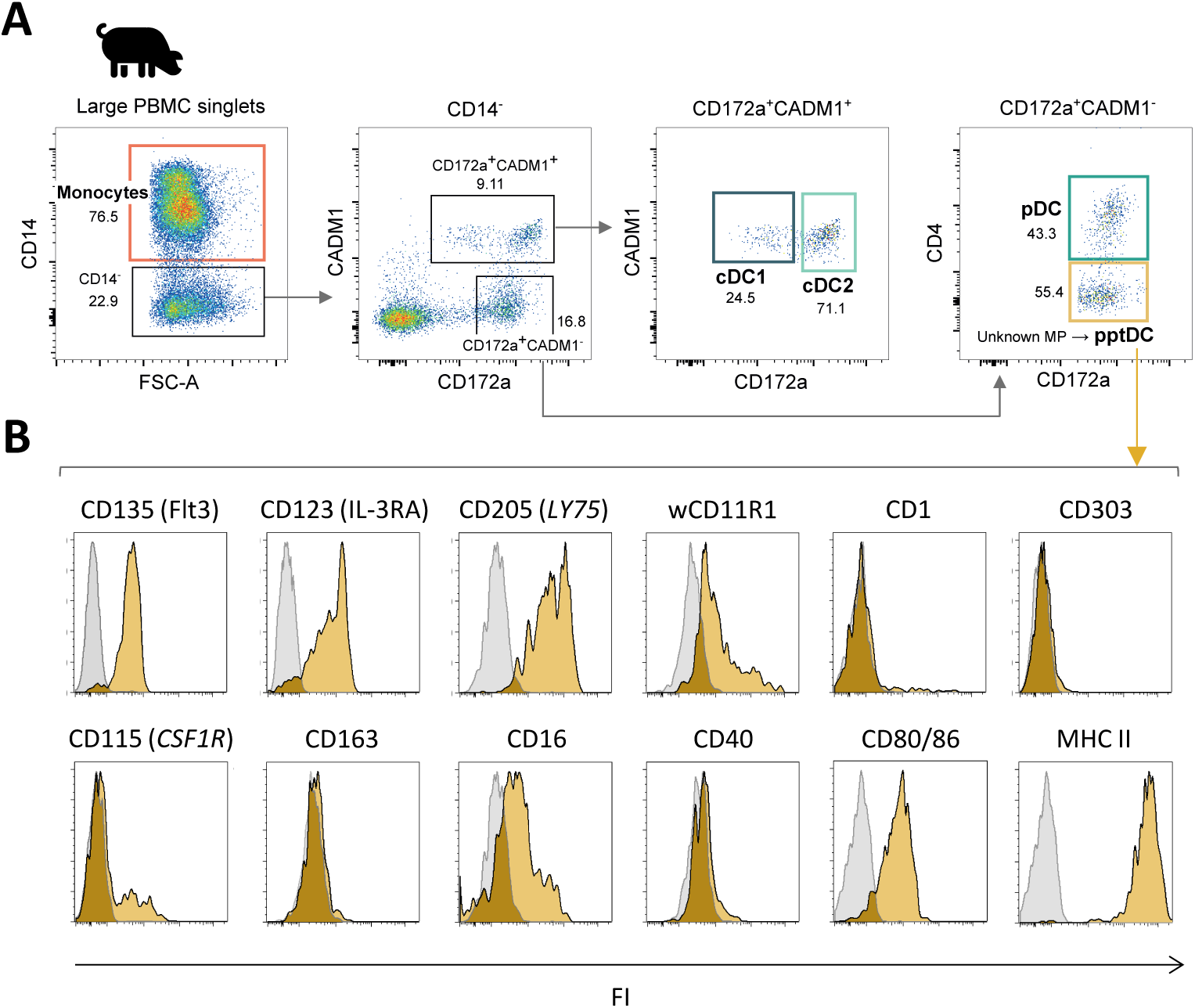
Phenotypic characterization of a new DC subset in blood of pigs: putative porcine tDC (pptDC). PBMC were isolated from the blood of four pigs and stained for flow cytometry. (A) Representative gating for mononuclear phagocyte (MP) subsets. Following selection of large cells and doublet exclusion (see **Supp. Fig. 1**), CD14^+^ cells were defined as monocytes and four subsets were distinguished among CD14-cells: cDC1 as CD172a^low^CADM1^+^, cDC2 as CD172a^+^CADM1^+^, pDC as CD172a^+^CADM1^-^CD4^+^, and a newly described DC subset as CD172a^+^CADM1^-^CD4^-^. (B) Yellow histograms show the expression of various molecules on CD172a^+^CADM1^-^CD4-cells. Gray histograms show the FMO control.

### Bulk transcriptome confirms identity of putative porcine tDC (pptDC)

Following phenotypic characterization, we investigated the transcriptional profile of the newly identified pptDC. To this end, this subset was MACS/FACS-sorted from the blood of four pigs and processed for bulk RNA-seq. Resulting data were analyzed alongside previously generated RNA-seq datasets of the four other blood MP subsets (cDC1, cDC2, pDC and monocytes) (5). Principal component analysis (PCA) of the 500 most variable genes between the five MP subsets (PC1 = 60%, PC2 = 30%) showed that all samples of the newly described cell subset (pptDC) were clustering together and away from the four other MP subsets, supporting the discovery of pptDC as a new and distinct DC subset (**Fig. 2A**). As further illustrated by a subset-to-subset correlation analysis, pptDC appeared to be more closely related to cDC than to pDC, sharing the highest correlation score with cDC2 (**Supp. Fig. 2A**).

**Figure 2:**
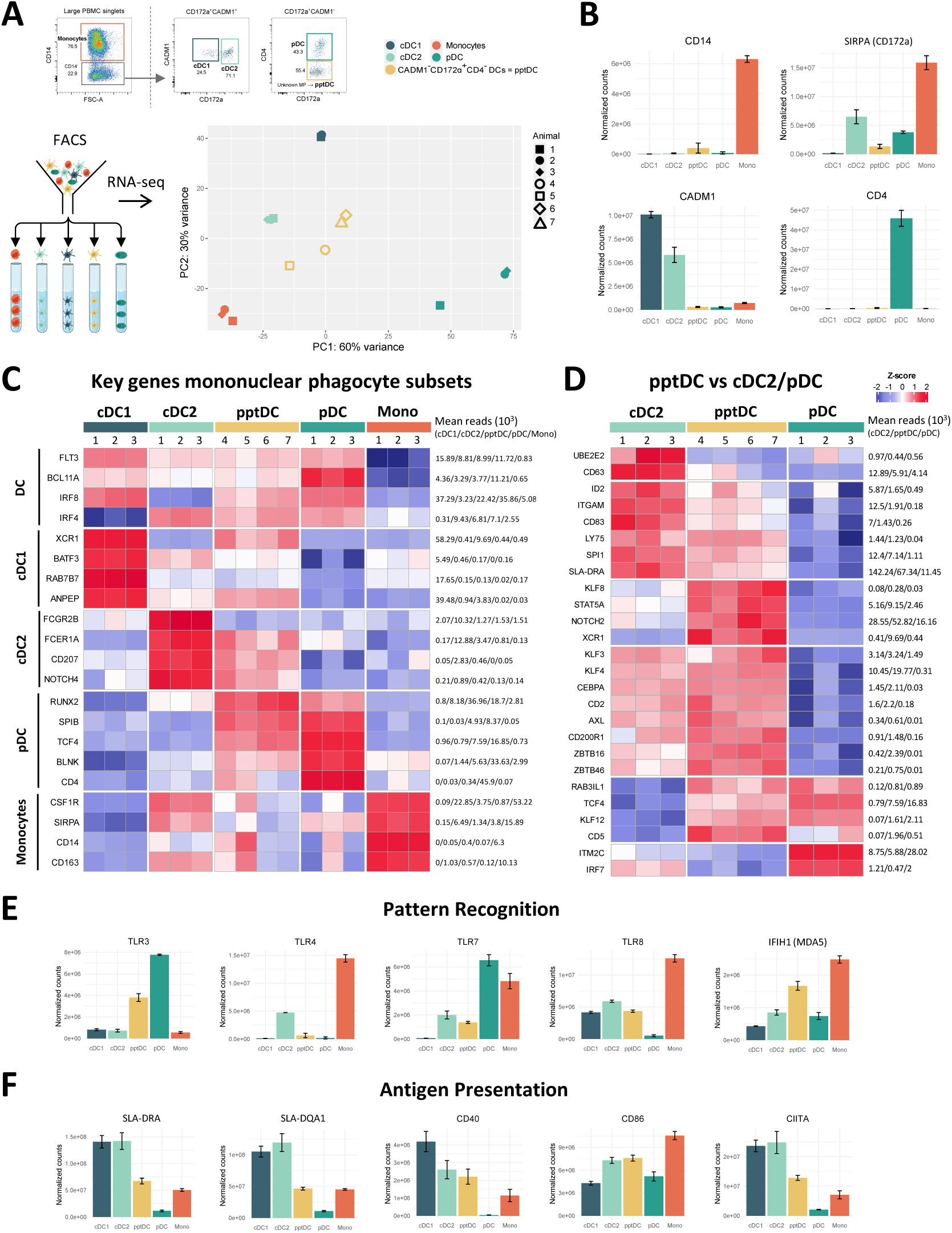
Bulk RNA-seq of DC subsets, including putative porcine tDC (pptDC) in blood of pigs. Bulk RNA-seq was performed on five sorted mononuclear phagocyte (MP) subsets. (**A**) First two dimensions of principal component analysis, with different symbols representing individual animals. (**B**) Gene expression for markers used in FACS. Bar plots show the number of normalized counts for each gene and MP subset (mean ± standard deviation). (**C**) Transcription of key MP subset-defining genes represented by heatmap. Z-scores were calculated from log10-transformed normalized counts of selected genes. Mean kilo reads for each subset and gene are given to the right of each heatmap. (**D**) Gene transcription distinguishing putative porcine tDC (pptDC) from cDC2 and/or pDC. Genes were selected based on pairwise comparisons with DESeq2 (adjusted p-value < 0.05 and |log2FC| > 1) (see **Supp. Data 1**) and literature research. (**E**-**F**) Gene transcription for pattern recognition and antigen presentation. Bar plots show the number of normalized counts for each gene and MP subset (mean ± standard deviation).

Overall, gene expression was in line with surface protein expression detected in flow cytometry (**Fig. 2B**). Gene expression for additional phenotypic markers is shown in **Supplementary Figure 2B**. Both pptDC and pDC expressed *IL3RA* (not available in Ensembl pig genome annotation), as shown by read mapping to the corresponding genomic region (**Supp. Fig. 2C**). To characterize pptDC, we next studied the expression of conserved key genes known to define the main MP subsets across species (53,54), including those we previously reported in pig blood (5). The updated analysis of the cDC1, cDC2, pDC and monocyte data (new reference genome) was in accordance with our formerly published transcriptomic analysis (5), supported also by the subset-specific expression pattern of these markers (**Fig. 2C**). For pptDC, the expression of the pan-DC markers *FLT3* and *BCL11A* was in common with cDC1, cDC2 and pDC. Considerable levels of monocyte-specific gene expression (e.g. *CSF1R*, *CD14*, *CD163*) were detected in two out of four pptDC samples (#5, #6 in **Fig. 2C**). Notably, pptDC expressed both IRF4- and IRF8-coding genes, two transcription factors (TF) involved in the development of cDC2 and cDC1/pDC, respectively (4,53). In addition to markers shared across DC populations, pptDC showed expression of DC subset-restricted features, such as *XCR1* and *ANPEP* (cDC1), *FCER1A*, *CD207* and *NOTCH4* (cDC2), and *RUNX2*, *TCF4*, *BLNK* and *SPIB* (pDC).

Dendritic cells with a phenotype and transcriptome overlapping with both cDC2 and pDC are characteristic of the recently identified tDC in humans (20) and mice (23). These tDC were reported to originate from progenitors shared with pDC and to differentiate into cDC2 (25). To further explore if our new DC subset represents the porcine equivalent of tDC, we analyzed differentially expressed genes (DEGs) between pptDC and cDC2 or pDC based on pairwise comparisons (DESeq2). Complete lists of DEGs are provided as **Supplementary Data 1**. Putative porcine tDC showed expression of *SPI1*, *TCF4*, *NOTCH2*, *CEPBA*, *KLF3*, *KLF8*, *KLF12*, *ZBTB46*, *IRF4*, *IRF8*, *STAT5A*, *RUNX2* and *SPIB* (**Fig. 2C-D**), which are genes also found in tDC of humans and mice (23,25,55,56), coding for cDC- or pDC-specific transcription factors (TF). Notably, Leylek *et al*. demonstrated by chromatin accessibility analysis that KLF3, KLF8 and KLF12 were part of the unique TF profile of tDC (55). Among the gene regulatory network governing DC development, TCF4 and ID2 are reported as mutual functional antagonists promoting pDC versus cDC differentiation, respectively (57). We found that both were expressed in pptDC (**Fig. 2D**), reinforcing its intermediate nature between cDC2 and pDC. Additionally, this new subset exhibited a high transcription level of *ZBTB16*, the gene encoding PLZF, a TF known to induce *ID2* expression (58). Interestingly, it shared the expression of *KLF4* with cDC2, reported to be required for the differentiation of murine circulating pDC-like cells (identified as pre-DC2) into a subset of cDC2 (27). Unlike pDC, pptDC showed low *IRF7* gene expression, suggesting its limited capacity to produce type I IFN, also distinguishing tDC from pDC in human and mouse (23). Moreover, its high transcription level of *CD200R1, STAT5A* and *RAB3IL1* in comparison to cDC2 and pDC is consistent with the spleen tDC signature described by Sulczewski *et al.* in mice (25).

Porcine cDC2 and pptDC could be further distinguished from pDC by their expression of *AXL*, a human tDC marker (**Fig. 2D**). Notably, pptDC expressed high levels of both *CD2* and *CD5*, in contrast to porcine pDC (low levels of *CD2* and *CD5* transcripts) and cDC2 (transcription of *CD2* but very low levels of *CD5* transcripts). These expression patterns were also found for human tDC, cDC2 and pDC, both transcriptionally and phenotypically (23). Thus, staining of CD2 and CD5 may be useful for distinction of porcine DC subsets in flow cytometry.

Finally, we observed a progressive increase of *ITGAM*, *CD83*, *LY75*, and *SLA-DRA* expression from pDC via pptDC toward cDC2 (**Fig. 2D**), suggesting gradually increasing antigen presentation capabilities across those subsets. Taken together, phenotype and bulk transcriptomic signatures support the idea that the new porcine DC subset (CD14^-^CADM1-CD172a^+^CD4^-^) represents the equivalent of tDC described in human and mouse.

### Functional specialization of pptDC inferred from bulk transcriptomics

Further exploration of pptDC-derived transcriptomic data revealed a unique gene signature related to pathogen recognition, antigen presentation, T-cell co-stimulation, immunoregulatory activities and cell adhesion (**Fig. 2** and **Supp. Fig. 3**).

Porcine putative tDC expressed relatively low levels of pattern recognition receptors (PPR) for bacterial components (e.g. *TLR4, TLR5*) (**Supp. Fig. 3A**), however one pptDC sample (#5) contained high transcript levels for bacterial PRR (**Supp. Fig. 3A**), and as shown in **Figure 2C**, also appeared enriched for monocyte-related transcripts such as *CD14*. Notably, all four pptDC samples stood out by high *TLR3* and *IFIH1* (*MDA-5*) expression (**Fig. 2E**), suggesting a specialization in sensing double stranded RNA. Transcripts for *TLR7*, *TLR8* and *TLR9* could also be detected in tDC, even though higher levels were detected in other DC subsets.

Similar to cDC2, pptDC expressed a relatively high level of C-type lectin receptor (CLR)-associated gene expression, such as *MRC1* (*CD206*), *PLA2R1 (CLEC13C)*, *CD207* and *CLEC4F* (**Supp. Fig. 3A**), suggesting their involvement in mannose-ligand recognition and phagocytic activities (59). Notably, expression of *MRC1* and *CLEC4F* was found to be very heterogeneous across pptDC samples. Similar to cDC1 and pDC, pptDC also contained transcripts for *CLEC12A* (*MICL*) and *CLEC12B*, two CLR that mostly recognize endogenous ligands such as damage-associated molecular patterns (60,61), thus suggesting a role in clearing dying cells. Looking at gene expression related to antigen presentation and T-cell modulation (**Supp. Fig. 3B**), pptDC stood out by expressing the highest levels of certain genes that may promote T-cell activation by enhancing antigen (cross-) presentation (*ATG5*, *UBE2D1, RAB27A)* (62–64), may promote the differentiation of regulatory T cells (*IL4I1*) (65,66) or Th1 cells (*DPP4*) (67), or may otherwise be involved in regulating T-cell responses (*VSIG10*, *CD200*) (68,69). For other genes involved in antigen presentation, we observed a gradual increase from pDC via pptDC towards cDC. Indeed, pptDC displayed intermediate transcription levels of genes encoding MHC-II molecules (e.g. *SLA-DRA*, *SLA-DOA*, *SLA-DMB*, *SLA-DMA*), molecules involved in MHC-II trafficking and antigen processing (*PIKFYVE*, *IFI30*, *CIITA*, *CD74*, *LY75*, *TAP2*) and co-stimulatory molecules (*CD83*, *CD40*) (**Fig. 2F, Supp. Fig. 2B** and 3B).

Genes encoding cytokines and cytokine receptors predominantly expressed in pptDC included *IL18* and *IL17RA* (**Supp. Fig. 3C**). Notably, alongside cDC1, pptDC prominently expressed the beta chain of the IL-6 receptor (IL6ST), reported to function in signal transduction for various cytokines (70). When compared to pDC, pptDC contained fewer transcripts for type I interferons (*IFN-OMEGA-6*) and related receptors (*IFNAR1* and *IFNAR2*), reinforcing the hypothesis of their limited involvement in type I IFN responses.

Looking at chemokines and chemokine receptors, pptDC were found to express considerable levels of *XCR1* (**Supp. Fig. 3D**). Notably, this key marker for cDC1 is involved in antigen cross-presentation and CD8^+^ T-cell priming (71),(72). Moreover, pptDC contained the highest number of *CCR7* and *CXCR5* transcripts among cDC subsets, however at low levels (mean reads of 200 and 300, respectively). While CCR7 is a well-known marker for DC activation (73), expression of CXCR5 is expected to cause migration to the CXCL13-rich parafollicular areas of the lymph node to possibly stimulate follicular Th cells (74).

Finally, several genes encoding integrin chains showed highest expression in pptDC such as *ITGB3*, *ITGAV* and *ITGA6* (**Supp. Fig. 3E**), as well as genes encoding Fc receptors (e.g. *FCRL4*) (**Supp. Fig. 3F**), metalloproteinases (e.g. *MME* and *MMP9*) (**Supp. Fig. 3G**) and semaphorins (e.g. *SEMA4F*, *SEMA4C*, *PLXNA4*) (**Supp. Fig. 3H**).

### Heterogeneity of porcine blood DC revealed by scRNA-seq

To get a more unbiased view on the heterogeneity of porcine blood DC subsets, we performed scRNA-seq (10x Genomics) on Flt3^+^ DC enriched from blood of three pigs (**Fig. 3A**). Clustering of cells with a resolution of 0.6 (Leiden algorithm) resulted in the identification of thirteen distinct clusters (**Fig. 3B**). Complete lists of cluster-defining marker genes, as determined by Seurat’s *FindAllMarkers()* function, are listed in **Supplementary Data 3**. Three clusters (c7, c8, c9) were excluded due to quality issues (**Supp. Fig. 4A**), as were clusters containing B cells (c13) and NK cells (c11) (**Supp. Fig. 4B-C**).

**Figure 3:**
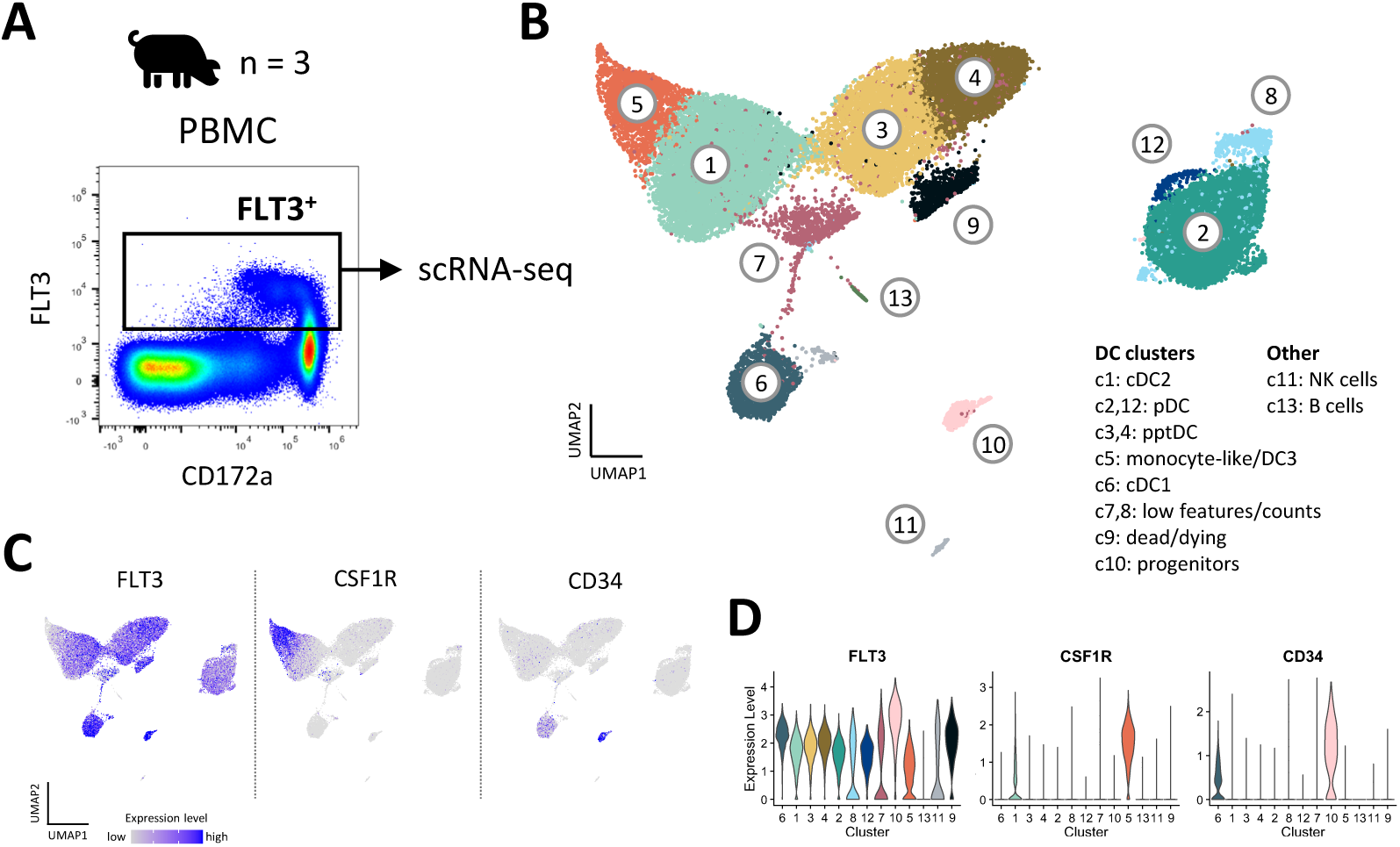
Heterogeneity of DC in blood of pigs revealed by scRNA-seq. (**A**) Dendritic cells (DC) were sorted from PBMC of three pigs and subjected to 10x Genomics scRNA-seq. Data from approximately 10 000 DC per sample was analyzed. (**B**) Clustering with a resolution of 0.6 resulted in 13 clusters visualized by UMAP plot. (**C**) Feature plots showing the expression of *FLT3, CSF1R* and *CD34*. (**D**) Violin plots showing the level of *FLT3, CSF1R and CD34* expression across all clusters.

Eight *FLT3*-expressing clusters (c1, c2, c3, c4, c5, c6, c10, and c12) were analyzed in further detail. Notably, cluster 5 appeared to contain monocytic cells alongside DC (DEG including *CSF1R*, *C5AR1*, *CD14*, *CD163*, *SIRPA* and *CD68*), and cluster 10 appeared to be comprised of DC progenitors (DEG including *CD34*, *MEIS1*, *DACH1*, *ERG, KIT*, *IKZF2* and *MECOM*) (16) (**Fig. 3C-D**, **Supp. Data 3**). As shown in **Figure 4A-C**, expression of subset-specific key genes clearly identified cluster 6 as cDC1 (*BATF3*, *XCR1*, *RAB7B*, *ANPEP, IRF8*), cluster 1 as cDC2 (*FCER1A*, *FCGR2B*, *CD207*, *SIRPA, IRF4*) and clusters 2 and 12 as pDC (*CD4*, *SPIB*, *BLNK*, *TCF4*, *RUNX2*, *IRF8*). Cells in clusters 3 and 4 expressed *IRF4*, *IRF8*, *XCR1*, *ANPEP*, *FCER1A*, *RUNX2*, *SPIB*, *TCF4* and *BLNK* (**Fig. 4B**), thus sharing subset-specific markers with cDC1, cDC2 and pDC, as observed with bulk RNA-seq of pptDC described above (**Fig. 2C**).

**Figure 4:**
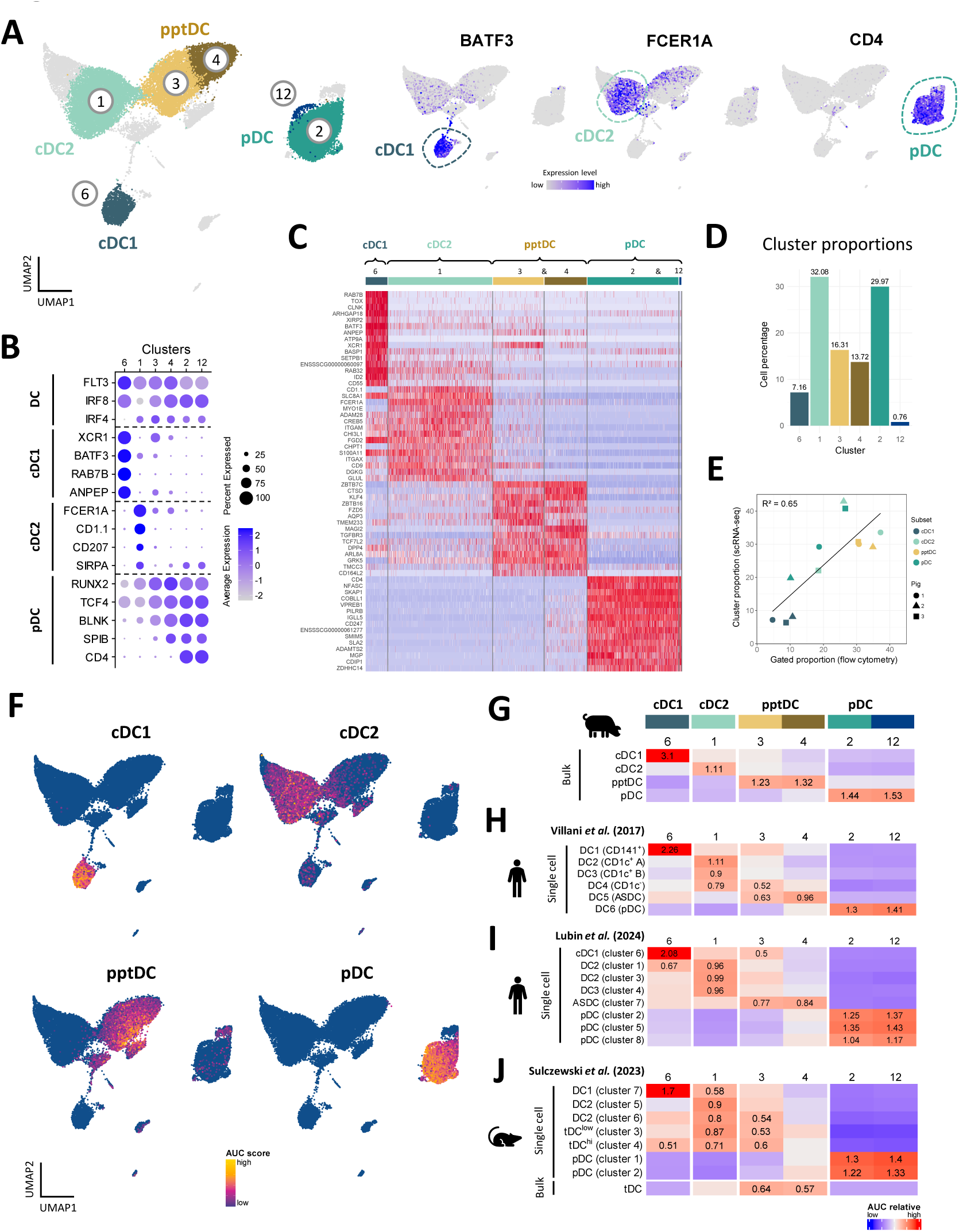
Annotation of DC subsets, including pptDC, in scRNA-seq dataset. (**A**) Subset annotation of main DC subsets (left) and expression of key genes visualized in feature plots (right). (**B**) Dot plot showing DC-subset-defining key genes. (**C**) Heatmap showing the top 15 differentially expressed genes (p_val_adj) for main DC clusters, as determined by Seurat’s *FindAllMarkers()* function. Complete gene lists are available as **Supp. Data 3**. (**D**) Proportion of each DC cluster relative to the total selected DC population. (**E**) Correlation between cell proportions obtained from scRNA-seq (y-axis) and flow cytometry (x-axis) of the same pigs. Symbols represent individual animals. Black line represents the linear regression model. (**F**) Enrichment of gene signatures from bulk-sequenced DC subsets (see **Supp. Data 1**) in single-cell RNA-seq clusters represented by UMAP plot. (**G**-**J**) Heatmaps showing averaged scaled enrichment scores for gene signatures from bulk-sequenced porcine DC subsets (see **Supp. Data 1**) (**G**), from human blood DC (Villani *et al*.) (**H**), from re-analyzed human blood DC (Lubin *et al*., for re-analysis see **Supp. Fig. 7**) (**I**), and from murine spleen DC (Sulczewski *et al*.) (**J**). AUC relative scores >= 0.5 are displayed.

A heatmap of the top 15 (adjusted p-value) DEGs between the DC subsets is shown in **Figure 4C** and the complete gene lists are given in **Supplementary Data 3**. Apart from the genes mentioned above, DEG included *CADM1*, *CLNK*, *ID2*, *SNX22, WDFY4* and *DNASEIL3*, for cDC1 (c6), *CD1D*, *ITGAM* (*CD11b*), *S100A4*, *TLR2* and *TLR4* for cDC2 (c1), and *IRF7*, *IFNAR1*, *NRP1*, *LRP8* and *SYK* for pDC (c2, c12). In line with the bulk RNA-seq analysis (**Fig. 2**, **Supp. Data 1**), clusters 3 and 4 were enriched for *ZBTB7C*, *ZBTB16*, *KLF4*, *NOTCH2*, *DPP4* (*CD26*), *TGFBR3* and *SEMA4F*, further supporting the classification of c3 and c4 as pptDC.

### Correlation of FCM-based and scRNA-seq based subset identification

When estimating the proportions of the DC clusters within total blood DC (**Fig. 4D**), we found that c1 (cDC2) and c2 (pDC) each represented approximately 30%, c6 (cDC1) represented 7%, and the pptDC clusters 3 and 4 represented 16% and 14%, respectively. These results are in accordance with the cell proportions previously observed by flow cytometry for the RNA-seq analysis of sorted cells (**Fig. 1A**). Representative flow cytometry data for animals included in the scRNA-seq analysis is shown in Supplementary Figure 5. Proportions of DC subsets from scRNA-seq and flow cytometry showed a moderate correlation, with an R² value of 0.65 (**Fig. 4E**).

To investigate if flow-cytometry defined DC subsets are well represented in the clustering of the scRNA-seq dataset, we tested for relative enrichment of the gene signatures from sorted bulk-sequenced subsets in the scRNA-seq clusters by gene-set enrichment analysis (GSEA). Different levels of enrichment were tested (5, 10, 15, 25, 50 and 100%) (see **Material and Methods**), allowing us to select 25% for optimal resolution of sc identities (**Supp. Fig. 6**). As shown in UMAP plots (**Fig. 4F**) and a heatmap (**Fig. 4G**), cDC1-derived gene sets had the highest enrichment in cluster 6, cDC2-derived gene sets in cluster 1, pDC-derived gene sets in clusters 2 and 12, and pptDC-derived gene sets in clusters 3 and 4. This clear allocation of bulk signatures supports the suitability of the 4-marker sorting strategy for porcine DC subsets and confirms the cluster identification in the scRNA-seq dataset.

### Porcine putative tDC share transcript signature with human and murine tDC

For further verification of pptDC identity, we used the GSEA approach described above to compare the porcine DC signatures to DC signatures derived from published bulk- and scRNA-seq studies of human blood (20,40) and murine spleen (25) (**Fig. 4H-J**). Two distinct human studies were selected, each utilizing a different scRNA-seq technology (SMARTSeq2 for Villani *et al.* (20), **Fig. 4H** and 10x Genomics for Lubin *et al.* (40), **Fig. 4I**). As expected, both human and murine cDC1, cDC2 and pDC gene signatures showed the highest relative enrichment in cluster 6 (cDC1), cluster 1 (cDC2) and in clusters 2 and 12 (both pDC), respectively. Notably, human ASDC and murine tDC (“bulk”) gene sets showed the highest enrichment in clusters 3 and 4, representing pptDC (**Fig. 4H-J**), thus supporting the identification of pptDC as porcine equivalents of tDC.

As expected, cluster 4 (pDC-like pptDC) was more enriched for human and murine pDC signatures than cluster 3 (cDC2-like pptDC), whereas cluster 3 displayed greater enrichment for human and murine cDC2 signatures than cluster 4. However, sc signatures of murine tDC-subclusters (“tDC^low^” and “tDC^hi^”) did not show discriminating enrichment in pptDC clusters 3 or 4 (**Fig. 4J**) but were rather enriched in c3 (cDC2-like pptDC) and c1 (cDC2).

### pptDC span a continuum between pDC-like and cDC-like profiles

In accordance with the reported origin of tDC from pro-pDC and their differentiation into cDC2-like cells (16,25,27,40,75), we found that transcripts for several TF involved in DC fate decisions (53) were sequentially increased or decreased from pDC via c4 and c3 towards cDC2 (**Fig. 5A**). Among TF, the most evident gradual decrease was observed for pDC-associated genes *TCF4*, *IRF8, IKZF1* and *BCL11A*, whereas transcription of cDC-associated genes *SPI1* and *ID2* increased via c4 and c3. A more abrupt decrease from c4 to c3 was observed for *SPIB*, the gene coding for Spi-B, a transcription factor promoting development of pDC (76). Notably, *RUNX2*, *NOTCH2*, *KLF4*, *KLF12* and *JAK2* showed highest expression in c4, before decreasing again in c3, whereas *ZBTB46* and *ZBTB16* expression appeared to peak in c3. Lastly, several TF showed increased expression in both c3 and c4 when compared to pDC and cDC2 (*ZBTB7C*, *STAT5A*, *KLF3* and *CBFA2T3*).

**Figure 5:**
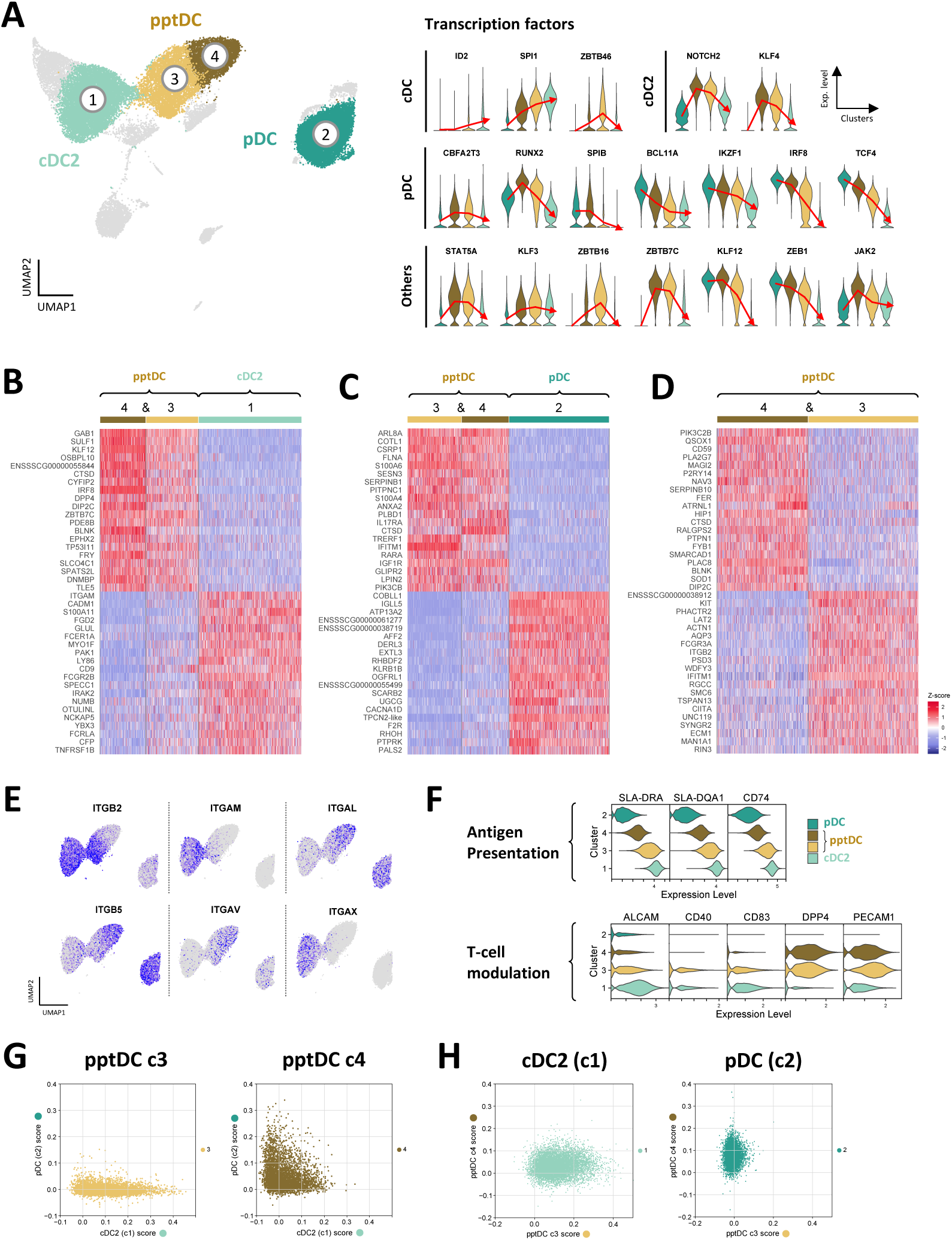
Transcriptomic delineation of pptDC from pDC and cDC2. Comparison of cDC2 (c1), pDC (focus on c2) and pptDC clusters (c3 and c4). (**A**) Violin plots show gene expression for transcription factors involved in the development of different DC subsets. Red lines pass through the mean expression value for each cluster. (**B-D**) Heatmaps show the top 20 differentially expressed genes (p_val_adj) in selected DC clusters, as determined by Seurat’s *FindMarkers()* function for pairwise comparisons. Complete gene lists are available as **Supp. Data 3**. (**E**) Feature plots showing the expression of genes coding for different integrin chains. **(F)** Expression of genes related to antigen presentation and T-cell modulation. **(G-H)** Single-cell classification scoring determined with an ElasticNet model.

The transcriptomic signature of pptDC clusters (c3 & c4) was further characterized by comparing them against cDC2 (c1) and pDC (c2) and against each other. Heatmaps of the top 20 (adjusted p-value) DEGs are shown in **Figure 5B-D**. The complete gene lists are given in **Supplementary Data 3**. Notably tDC clusters differed in the expression of integrin transcripts (**Fig. 5E**), generally reflecting their similarity to either pDC (c4: *ITGAL*, *ITGB5*) or cDC2 (c3: *ITGB2*, *ITGAM*, *ITGAX*). In accordance with the adoption of an increasingly cDC-like phenotype, cDC2-like tDC (c3) showed higher expression of transcripts related to antigen presentation (e.g. *SLA-DRA*, *SLA-DQA1*, *CD74*) and T-cell stimulation (e.g. *CD40*, *CD83*, *ALCAM*) than pDC-like tDC (c4) (**Fig. 5G**). In line with bulk RNA-seq, pptDC clusters showed the highest expression of *DPP4* and *PECAM1* (two genes involved in T-cell modulation), when compared to pDC and cDC2.

To further characterize the correlation between pptDC and cDC2 or pDC, we trained an ElasticNet model computing a classification score based on the transcriptomic signatures of the different cell types (**Fig. 5G-H**). According to this model, pptDC in cluster 4 showed low-to-intermediate scores for both cDC2 and pDC signatures, while pptDC in cluster 3 were characterized by a progressive increase in the cDC2 score while exhibiting a low pDC score (**Fig. 5G**). In line with the differential gene expression analysis, cDC2 displayed a higher score for cluster 3 than for cluster 4, and pDC showed the reverse pattern (**Fig. 5H**).

Overall, the transcriptome of pptDC spans a continuum between pDC and cDC profiles, as described for humans and mice (20,23,25,77). This transient transcriptome, together with the core gene signature resembling human and murine tDC, justifies identifying pptDC as the porcine equivalents of ASDC/tDC. Furthermore, these results suggest that porcine tDC may differentiate into cDC2-like cells, as proposed for their murine (25,27) and human counterparts (40).

To further explore the cellular dynamics between porcine tDC and cDC2, we performed a trajectory inference (TI) analysis on cells in clusters 1, 3 and 4 (**Supp. Fig. 8**). Genes that varied along the inferred trajectory corresponded to the DC subset-specific signatures previously identified by differential expression analysis. Notably, the continuum from tDC to cDC2 was marked by a gradual increase in the expression of genes associated with antigen presentation via MHC-II molecules (module 2 in **Supp. Fig. 8C**), suggesting a progressive acquisition of cDC features by tDC differentiating towards cDC2-like cells. Results of

### Putative DC3 in porcine blood

In both mouse and human, DC3 have been described as a novel DC subset sharing dendritic and monocytic markers (13,15,18–20). Interestingly, cluster 5, adjacent to cDC2 (c1), appeared to contain both monocytic and dendritic cells (**Fig. 3**) and showed the highest enrichment score when performing GSEA with human DC3 signatures (**Fig. 6A** and **6B**).

**Figure 6:**
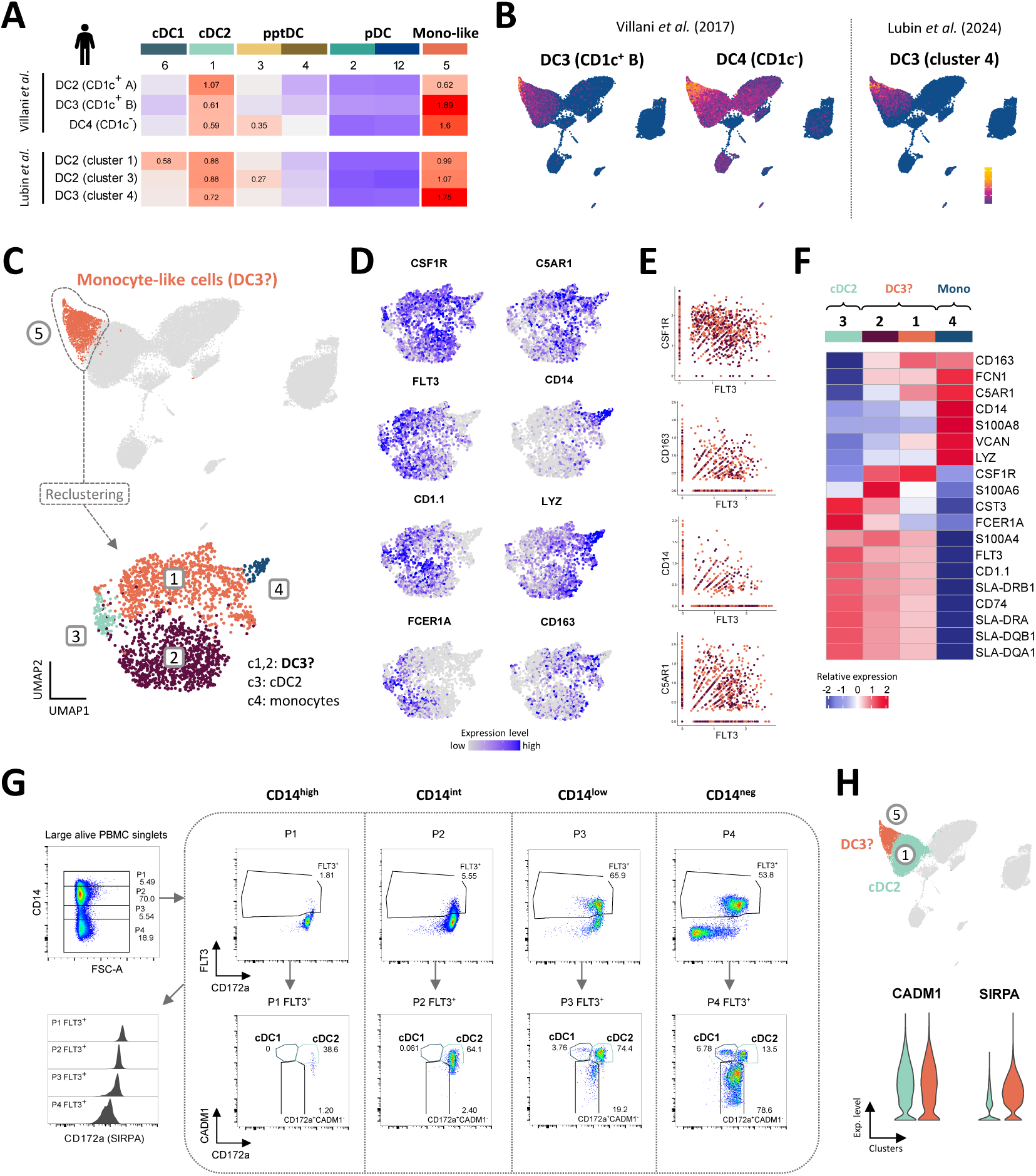
Putative porcine DC3 co-express Flt3 and CD14. (A-B) Enrichment of gene signatures from human blood DC2 and DC3 (Villani *et al.* and Lubin *et al.*), represented by heatmaps(A) and UMAP plots (B). AUC relative scores >= 0.5 are displayed. (C) Re-clustering of monocyte-like cells in cluster 5, resulting in 4 distinct clusters visualized by UMAP plot (Leiden algorithm, clustering resolution = 0.4). (D) Feature plots show expression of selected genes reported to distinguish human cDC2, DC3 and monocytes. (E) Scatter plots show co-expression of *FLT3* and selected monocytic markers in clusters 1 and 2. (F) Heatmap shows relative mean gene expression across each cluster for selected genes reported to distinguish between human cDC2, DC3 and monocytes. (G) Flow-cytometric gating of porcine DC subsets from gradual CD14 expression gates (P1-4). Plots are representative for all three pigs included in the scRNA-seq analysis. For complete gating strategy see Supp. Fig. 5. (H) Violin plots showing the expression of genes encoding CADM1 and CD172a (*SIRPA*) for the cDC2 (c1) and DC3 clusters (c5) in the original scRNA-seq dataset.

Upon re-clustering of c5, two major clusters were separated from two smaller clusters (**Fig. 6C**). The small clusters 3 and 4 were annotated as cDC2 (*FLT3*, *FCER1A*, *CD1.1*) and monocytes (high levels of *CD14*, *LYZ*), respectively, while the two major clusters 1 and 2 were annotated as putative DC3 as they contained low levels of *CD14* and *FCER1A* transcripts alongside *FLT3* and *CSF1R* transcripts (**Fig. 6C** and **6D**). Indeed, a significant proportion of cells in sub-clusters 1 and 2 (putative DC3) were revealed to co-express *FLT3* transcripts alongside transcripts typically associated with monocytes, such as *CSF1R*, *CD163, CD14* and *C5AR1* (**Fig. 6E**).

Among putative DC3, c1 stood out by higher transcription of *CD163*, *C5AR1* and *VCAN*, while c2 was clearly enriched in *S100A4*, *S100A6* and *CST3* transcripts (**Fig. 6F**). Compared to c1, c2 also expressed slightly higher levels of MHC-II-related genes.

The current gating strategy for identifying porcine DC subsets is based on exclusion of CD14^+^ monocytes (**Fig. 1**) and therefore likely excludes CD14-expressing DC3. This is illustrated by gradual gating based on CD14 expression prior to DC gates, as shown in **Figure 6G**. Indeed, CD14^low^ (P3) and CD14^int^ (P2) populations contained approximately 66% and 6% Flt3^+^ DC, respectively, mainly falling within the cDC2 gate (CADM1^+^CD172a^+^). The gated Flt3^+^CADM1^+^CD172a^+^ population thus represents a more heterogenous population, likely containing CD14-expressing DC3. This hypothesis is further supported by the shared expression of CADM1 and CD172a (*SIRPA*) by cDC2 (c3) and putative DC3 (c5) at the transcriptomic level (**Fig. 6H**).

## DISCUSSION

We have previously identified porcine cDC1, cDC2 and pDC in blood of pigs by their expression of key transcripts conserved across species (5). In this previous work, we found a substantial subset of CD14^-^CADM1^-^CD172a^+^CD4- cells in the blood of pig with unknown identity (5). Based on recent insights from human and mouse, the present study now identifies this unknown DC subset as the equivalent of tDC by combining flow cytometry, bulk- and scRNA-seq analyses. With the current study, we have zoomed into the DC compartment by performing scRNA-seq on Flt3-enriched PBMC, revealing both tDC and putative DC3 in blood of pigs. Like their human and murine counterparts (20,23,25,78), porcine tDC displayed a distinct transcriptomic signature in-between pDC and cDC2, whereas putative DC3 clustered in a continuum in-between cDC2 and monocytes.

Notably, porcine tDC were found to be as frequent as other DC subsets in blood of pigs, which is in stark contrast to reports from human and mouse, where tDC only form a minor population of approximately 1-5 % among total DC in blood and spleen (20,23,25,27,40,77). The high proportion of tDC in porcine blood is puzzling and may point towards high frequencies of tDC across tissues, which would make the pig an attractive model for studying tDC in various settings, including infection. In fact, tDC are discussed to play a special role in viral infection. In murine models of SARS-CoV2 infection, virus-sensing tDC produced IL1-β and were deemed responsible for shifting the balance towards inflammation and fatal immunopathology (25). Upregulation of IL1-β was also observed in human tDC recruited to skin following experimental injection of UV-killed *E. coli* (40). Compared to blood tDC, these tDC had upregulated pro-inflammatory genes (*IL1B, SAT1, AXL*), genes related to IFN signaling (*ISG15*, *IFI44L*, *IFI27*), and surface chemokine receptors associated to migration (CXCR4, CX3CR1), while having downregulated genes coding for HLA molecules.

Our transcriptomic data support the involvement of porcine tDC in sensing viral components and in promoting inflammation. In particular, sensing double-stranded RNA via TLR3 seems to be important in porcine tDC. Notably, apart from porcine tDC, TLR3 is predominantly expressed in porcine pDC. This is in contrast to human and mice, where TLR3 is not expressed at all on pDC (79,80), representing only one example of species differences in viral sensing (5).

Porcine tDC expressed higher levels of CD86 and MHC-II molecules than pDC, both on the mRNA and protein level, suggesting that tDC are better equipped for T-cell stimulation. This is in line with murine and human tDC reported to outperform pDC in inducing allogenic T-cell proliferation (20,23,25,27,40). In fact, contaminations with tDC/human ASDC in traditional pDC gates have likely biased T-cell stimulation assays, erroneously attributing T-cell stimulatory functions to pDC (20). It remains to be determined if tDC contribute to stimulation of naive T cells in secondary lymphoid tissues. In a model of murine influenza infection, tDC were described to be recruited to the lungs, but were not found to accumulate in draining lymph nodes (23). While our data indicate transcription of several TLR and co-stimulatory molecules, future studies should interrogate TLR responsiveness and the capacity of porcine tDC for phenotypic maturation (upregulation of CCR7, MHC-II, CD80/86), as previously performed for bovine DC and monocyte subsets (9,81). In particular, assessment of CCR7 upregulation upon TLR stimulation will indicate if porcine tDC are capable of migration to T-cell zones in secondary lymphoid tissues. The transcriptomic reference datasets generated in the present study will enable detailed investigations on tDC and their activation signatures across tissues both in steady-state and infection.

Murine studies have started to dissect the developmental pathway of tDC using specific knockout (KO) and adoptive cell transfer approaches, as well as lineage tracing mouse models (16,25,27). Sulczewski *et al.* demonstrated that murine tDC originate from bone marrow progenitors shared with pDC (pro-pDC) at steady state (25). Notably, when knocking out the pre-cDC pathway, pro-pDC could compensate for the lack of cDC2 by producing cDC2-like cells (termed tDC2) via the tDC pathway. Moreover, tDC isolated from human blood converted into CD5^+^ cDC2 upon CD40L stimulation *in vitro* (25) and bone marrow tDC cultured under standard DC differentiation conditions (i.e. GM-CSF and FLT3-ligand) generated exclusively DC2 (75). The clustering we observed in our scRNA-seq dataset alongside the transiently increasing cDC signature would support the hypothesis that porcine tDC can give rise to cDC2-like cells *in vivo*. It is intriguing to speculate that the bridge-like connection between cDC2-like tDC and cDC2 in our UMAP plot marks this transition. Recently, Lubin *et al*. used deuterium-glucose labeling of dividing cells to study the kinetics of DC subsets and their progenitors in human blood (40). Their findings support a model in which ASDC (human tDC) give rise to DC2 in both the bone marrow and blood. Indeed, the incorporation of deuterium into DNA is safe for use in humans and rodents (82,83), making it a promising tool for investigating the fate and lifespan of DC in large animal models *in vivo*.

Patterns of expressed transcription factors in porcine tDC were largely in accordance with murine and human tDC. High expression of *STAT5A* in porcine tDC is in line with the idea that tDC need to counteract differentiation towards pDC, as STAT5 was reported to inhibit pDC development by suppressing IRF8 (84). By chromatin accessibility analysis, Leylek *et al*. demonstrated that KLF3, KLF8 and KLF12 were part of the unique TF profile of tDC (55). Accordingly, the distinct expression pattern of *KLF3*, *KLF8* and *KLF12* in porcine tDC distinguished them from cDC2 and pDC. Among the gene regulatory network governing DC development, TCF4 and ID2 are reported as mutual functional antagonists promoting pDC versus cDC differentiation, respectively (57). The expression of both *TCF4* and *ID2* in porcine tDC aligns with their transitional nature.

To our knowledge, tDC have not yet been described in mammalian species other than humans and mice. A decade ago, Vu Manh *et al.* described a subpopulation of cDC2 (FSC^hi^MHC-II^+^CD14^-^CD4^-^CADM1^-^CD172a^int^) in porcine blood (54). The bulk transcriptome of this population was suggested to be significantly contaminated by pDC (*TCF4*) and cDC1 (*XCR1*) and was thus excluded from their analyses. In the light of current knowledge and our present results, this population likely contained tDC. As did the *FLT3-* and *XCR1*-expressing CADM1-population within CD14^-^CD172a^+^CD1^-^CD4-cDC reported in another study (85).

Transcriptomic data suggest that CD2 and CD5 can be used in flow cytometry to discriminate porcine tDC (CD2^+^CD5^+^) from pDC (CD2^low^CD5^low^) and cDC2 (CD2^+^CD5^low^). Notably, human tDC, previously considered as pre-DC, are also reported to differ from pDC by expression of CD2 and CD5 transcripts (20,78). Protein-level analyses are necessary to confirm the suitability of these markers. The combination of bulk RNA-seq from sorted DC populations and scRNA-seq of enriched DC allowed us to confirm the accuracy of our flow-cytometry based subset identification (cDC1, cDC2, pDC, tDC). However, scRNA-seq of enriched DC revealed additional heterogeneity. Unbiased clustering of our scRNA-seq dataset suggests the presence of two tDC subsets in porcine blood, spanning a differentiation continuum in-between pDC-like cells and cDC2-like cells. Similarly, in mice, tDC were classified into two distinct subpopulations according to their similarity to pDC and cDC2, termed e tDC^low^ (CD11c^low^Ly6c^high^) and tDC^high^ (CD11c^high^Ly6C^low^), respectively (23). The mouse *Ly6c* gene does not have a pig ortholog, but transcription of *ITGAX*, encoding CD11c, appeared to be higher in porcine cDC2-like tDC. So, although transcriptomic signatures from murine tDC^high^ and tDC^low^ were not discriminatory for the two porcine tDC clusters, CD11c may still be suitable for distinguishing porcine tDC subsets in flow cytometry. Observed monocyte signatures (increased transcripts for e.g. *CD14*) in two out of four investigated tDC samples are surprising and cannot be explained by the scRNA-seq data, where monocyte-associated gene expression could not be detected in the two tDC clusters.

In addition to tDC, our scRNA-seq analyses suggest the presence of DC3 in porcine blood. Dendritic cells type 3 have been described as a new DC lineage, originating from monocyte/dendritic cell precursors, as opposed to cDC deriving from common dendritic progenitors (16,19). As DC3 share phenotype and transcriptome with monocytes and cDC2, their clear delineation has proven difficult in both human and mouse, especially under inflammatory conditions (86). In fact, CD14, a molecule that has traditionally been used as a monocyte marker across species, appears to be expressed on DC3 of all species investigated so far, including pig. Notably, in the gating strategy employed here to sort porcine DC subsets for bulk RNA-seq, CD14^+^ cells were excluded. This has likely reduced DC3 contamination in the cDC2 gate, but also highlights the importance of scRNA-seq as a tool that is relatively independent of *a priori* defined gating strategies. In future studies, the gating strategy for porcine DC subsets should include Flt3 to account for CD14-expressing DC3.

When studying rare and poorly defined DC with scRNA-seq, proper enrichment strategies are crucial and should be based on extensive phenotypic characterization to not bias investigations on DC heterogeneity. By enriching DC by Flt3 expression, as performed in the present study, we expect to have captured the vast majority of DC. In support of this, similar proportions for main DC subsets were found in scRNA-seq (Flt3-enriched) and flow cytometry (Flt3-independent gating strategy employed for bulk RNA-seq of DC subsets). However, DC subsets expressing low levels of Flt3 (e.g. pDC) may still be missed by this enrichment strategy, as also discussed for scRNA-seq of Flt3-enriched cells in bovine lymph node (87).

Taken together, by enriching Flt3^+^ cells for scRNA-seq, we have zoomed into the heterogeneous compartment of porcine DC at unprecedented detail. Apart from discovering tDC as a major DC subset in porcine blood, we describe putative DC3 as FLT3 expressing cells that show considerable transcriptional overlap with monocytes. Several open questions need to be addressed in future studies, including the functional role of these DC subsets across species, and the suitability of the pig as a model species for human tDC research.

## DATA AVAILABILITY STATEMENT

Raw sequencing data from bulk RNA-seq of pig blood cDC1, cDC2, pDC and monocytes are available in the European Nucleotide Archive (ENA) (http://www.ebi.ac.uk/ena) under the accession number PRJEB15381. Raw sequencing data from bulk RNA-seq of porcine blood tDC and from scRNA-seq will be available in ENA upon publication of the final version.

## CODE AVAILABILITY

Scripts used for read alignments to the pig reference genome and the bulk and scRNA-seq data analyses will be available in a GitHub public repository upon publication of the final version.

## AUTHOR CONTRIBUTIONS

AB performed bioinformatics data analysis and prepared figures and text. GA performed laboratory work and data analysis. AH performed laboratory work and analyzed the flow cytometry data. MB provided support with bioinformatics data analysis and laboratory work. FB supervised bioinformatics data analysis. AS participated in the design of the study, supervised data analysis and edited the manuscript. SCT participated in the design of the study, supervised laboratory work and data analysis and edited the manuscript.

## ACKNOWLEDGEMENTS

We thank Sylvie Python and Caroline Lehmann (IVI, Bern, Switzerland) for their support in the laboratory, animal caretakers at IVI for blood sampling, Stefan Müller (FCCS, University of Bern, Switzerland) for cell sorting, Pamela Nicholson (NGS Platform, University of Bern, Switzerland) for single-cell RNA sequencing, and the Interfaculty Bioinformatics Unit (IBU, University of Bern, Switzerland) for access to their compute cluster.

## REFERENCES

1. Cabeza-Cabrerizo, M., Cardoso, A., Minutti, C. M., Pereira Da Costa, M. & Reis E Sousa, C. Dendritic Cells Revisited. Annu. Rev. Immunol. 39, 131–166 (2021).1. Villar J, Segura E. Decoding the Heterogeneity of Human Dendritic Cell Subsets. Trends in Immunology. 2020 Dec;41(12):1062–71.

2. Collin M, Bigley V. Human dendritic cell subsets: an update. Immunology. 2018 May;154(1):3–20.

3. Cabeza-Cabrerizo M, Cardoso A, Minutti CM, Pereira Da Costa M, Reis E Sousa C. Dendritic Cells Revisited. Annu Rev Immunol. 2021 Apr 26;39(1):131–66.

4. Zhang S, Audiger C, Chopin M, Nutt SL. Transcriptional regulation of dendritic cell development and function. Front Immunol. 2023 Jul 14;14:1182553.

5. Auray G, Keller I, Python S, Gerber M, Bruggmann R, Ruggli N, et al. Characterization and Transcriptomic Analysis of Porcine Blood Conventional and Plasmacytoid Dendritic Cells Reveals Striking Species-Specific Differences. The Journal of Immunology. 2016 Dec 15;197(12):4791–806.

6. Auray G, Talker SC, Keller I, Python S, Gerber M, Liniger M, et al. High-Resolution Profiling of Innate Immune Responses by Porcine Dendritic Cell Subsets in vitro and in vivo. Front Immunol. 2020 Jul 7;11:1429.

7. Summerfield A, Auray G, Ricklin M. Comparative dendritic cell biology of veterinary mammals. Annu Rev Anim Biosci. 2015;3:533–57.

8. Talker SC, Baumann A, Barut GT, Keller I, Bruggmann R, Summerfield A. Precise Delineation and Transcriptional Characterization of Bovine Blood Dendritic-Cell and Monocyte Subsets. Front Immunol. 2018 Oct 30;9:2505.

9. Barut GT, Lischer HEL, Bruggmann R, Summerfield A, Talker SC. Transcriptomic profiling of bovine blood dendritic cells and monocytes following TLR stimulation. Eur J Immunol. 2020 Nov;50(11):1691–711.

10. Ziegler A, Marti E, Summerfield A, Baumann A. Identification and characterization of equine blood plasmacytoid dendritic cells. Developmental & Comparative Immunology. 2016 Dec;65:352–7.

11. Baillou A, Tomal F, Chaumeil T, Barc C, Levern Y, Sausset A, et al. Characterization of intestinal mononuclear phagocyte subsets in young ruminants at homeostasis and during Cryptosporidium parvum infection. Front Immunol. 2024 May 2;15:1379798.

12. Chen B, Zhu L, Yang S, Su W. Unraveling the Heterogeneity and Ontogeny of Dendritic Cells Using Single-Cell RNA Sequencing. Front Immunol. 2021 Sep 9;12:711329.

13. Dutertre CA, Becht E, Irac SE, Khalilnezhad A, Narang V, Khalilnezhad S, et al. Single-Cell Analysis of Human Mononuclear Phagocytes Reveals Subset-Defining Markers and Identifies Circulating Inflammatory Dendritic Cells. Immunity. 2019 Sep;51(3):573–589.e8.

14. Minutti CM, Piot C, Pereira Da Costa M, Chakravarty P, Rogers N, Huerga Encabo H, et al. Distinct ontogenetic lineages dictate cDC2 heterogeneity. Nat Immunol. 2024 Mar;25(3):448–61.

15. Cytlak U, Resteu A, Pagan S, Green K, Milne P, Maisuria S, et al. Differential IRF8 Transcription Factor Requirement Defines Two Pathways of Dendritic Cell Development in Humans. Immunity. 2020 Aug;53(2):353–370.e8.

16. Rodrigues PF, Trsan T, Cvijetic G, Khantakova D, Panda SK, Liu Z, et al. Progenitors of distinct lineages shape the diversity of mature type 2 conventional dendritic cells. Immunity. 2024 Jul;57(7):1567–1585.e5.

17. León B. Type 2 conventional dendritic cell functional heterogeneity: ontogenically committed or environmentally plastic? Trends in Immunology. 2025 Jan;S1471490624003077.

18. Bourdely P, Anselmi G, Vaivode K, Ramos RN, Missolo-Koussou Y, Hidalgo S, et al. Transcriptional and Functional Analysis of CD1c+ Human Dendritic Cells Identifies a CD163+ Subset Priming CD8+CD103+ T Cells. Immunity. 2020 Aug;53(2):335–352.e8.

19. Liu Z, Wang H, Li Z, Dress RJ, Zhu Y, Zhang S, et al. Dendritic cell type 3 arises from Ly6C+ monocyte-dendritic cell progenitors. Immunity. 2023 Aug;56(8):1761–1777.e6.

20. Villani AC, Satija R, Reynolds G, Sarkizova S, Shekhar K, Fletcher J, et al. Single-cell RNA-seq reveals new types of human blood dendritic cells, monocytes, and progenitors. Science. 2017 Apr 21;356(6335):eaah4573.

21. Rodrigues PF, Alberti-Servera L, Eremin A, Grajales-Reyes GE, Ivanek R, Tussiwand R. Distinct progenitor lineages contribute to the heterogeneity of plasmacytoid dendritic cells. Nat Immunol. 2018 Jul;19(7):711–22.

22. Arroyo Hornero R, Idoyaga J. Plasmacytoid dendritic cells: A dendritic cell in disguise. Molecular Immunology. 2023 Jul;159:38–45.

23. Leylek R, Alcántara-Hernández M, Lanzar Z, Lüdtke A, Perez OA, Reizis B, et al. Integrated Cross-Species Analysis Identifies a Conserved Transitional Dendritic Cell Population. Cell Reports. 2019 Dec;29(11):3736–3750.e8.

24. Matsui T, Connolly JE, Michnevitz M, Chaussabel D, Yu CI, Glaser C, et al. CD2 Distinguishes Two Subsets of Human Plasmacytoid Dendritic Cells with Distinct Phenotype and Functions. The Journal of Immunology. 2009 Jun 1;182(11):6815–23.

25. Sulczewski FB, Maqueda-Alfaro RA, Alcántara-Hernández M, Perez OA, Saravanan S, Yun TJ, et al. Transitional dendritic cells are distinct from conventional DC2 precursors and mediate proinflammatory antiviral responses. Nat Immunol. 2023 Aug;24(8):1265–80.

26. Salvermoser J, Van Blijswijk J, Papaioannou NE, Rambichler S, Pasztoi M, Pakalniškytė D, et al. Clec9a-Mediated Ablation of Conventional Dendritic Cells Suggests a Lymphoid Path to Generating Dendritic Cells In Vivo. Front Immunol. 2018 Apr 16;9:699.

27. Rodrigues PF, Kouklas A, Cvijetic G, Bouladoux N, Mitrovic M, Desai JV, et al. pDC-like cells are pre-DC2 and require KLF4 to control homeostatic CD4 T cells. Sci Immunol. 2023 Feb 3;8(80):eadd4132.

28. Radulovic E, Mehinagic K, Wüthrich T, Hilty M, Posthaus H, Summerfield A, et al. The baseline immunological and hygienic status of pigs impact disease severity of African swine fever. Dixon LK, editor. PLoS Pathog. 2022 Aug 25;18(8):e1010522.

29. Bolger AM, Lohse M, Usadel B. Trimmomatic: a flexible trimmer for Illumina sequence data. Bioinformatics. 2014 Aug 1;30(15):2114–20.

30. Dobin A, Davis CA, Schlesinger F, Drenkow J, Zaleski C, Jha S, et al. STAR: ultrafast universal RNA-seq aligner. Bioinformatics. 2013 Jan 1;29(1):15–21.

31. Liao Y, Smyth GK, Shi W. featureCounts: an efficient general purpose program for assigning sequence reads to genomic features. Bioinformatics. 2014 Apr 1;30(7):923–30.

32. Love MI, Huber W, Anders S. Moderated estimation of fold change and dispersion for RNA-seq data with DESeq2. Genome Biol. 2014 Dec;15(12):550.

33. R Core Team. R: A Language and Environment for Statistical Computing [Internet]. Vienna, Austria: R Foundation for Statistical Computing; 2023. Available from: https://www.R-project.org/

34. Zheng GXY, Terry JM, Belgrader P, Ryvkin P, Bent ZW, Wilson R, et al. Massively parallel digital transcriptional profiling of single cells. Nat Commun. 2017 Jan 16;8(1):14049.

35. Hao Y, Stuart T, Kowalski MH, Choudhary S, Hoffman P, Hartman A, et al. Dictionary learning for integrative, multimodal and scalable single-cell analysis. Nat Biotechnol [Internet]. 2023 May 25 [cited 2024 Jul 22]; Available from: https://www.nature.com/articles/s41587-023-01767-y

36. Germain PL, Lun A, Garcia Meixide C, Macnair W, Robinson MD. Doublet identification in single-cell sequencing data using scDblFinder. F1000Res. 2022 May 16;10:979.

37. Korsunsky I, Fan J, Slowikowski K, Zhang F, Wei K, Baglaenko Y, et al. Fast, sensitive, and accurate integration of single cell data with Harmony [Internet]. 2018 [cited 2024 Jul 23]. Available from: http://biorxiv.org/lookup/doi/10.1101/461954

38. Aibar S, González-Blas CB, Moerman T, Huynh-Thu VA, Imrichova H, Hulselmans G, et al. SCENIC: single-cell regulatory network inference and clustering. Nat Methods. 2017 Nov;14(11):1083–6.

39. Herrera-Uribe J, Wiarda JE, Sivasankaran SK, Daharsh L, Liu H, Byrne KA, et al. Reference Transcriptomes of Porcine Peripheral Immune Cells Created Through Bulk and Single-Cell RNA Sequencing. Front Genet. 2021 Jun 23;12:689406.

40. Lubin R, Patel AA, Mackerodt J, Zhang Y, Gvili R, Mulder K, et al. The lifespan and kinetics of human dendritic cell subsets and their precursors in health and inflammation. Journal of Experimental Medicine. 2024 Nov 4;221(11):e20220867.

41. Pedregosa F, Varoquaux G, Gramfort A, Michel V, Thirion B, Grisel O, et al. Scikit-learn: Machine Learning in Python. Journal of Machine Learning Research. 2011;12:2825–30.

42. Mayère C, Regard V, Perea-Gomez A, Bunce C, Neirijnck Y, Djari C, et al. Origin, specification and differentiation of a rare supporting-like lineage in the developing mouse gonad. Sci Adv. 2022 May 27;8(21):eabm0972.

43. Zou H, Hastie T. Regularization and Variable Selection Via the Elastic Net. Journal of the Royal Statistical Society Series B: Statistical Methodology. 2005 Apr 1;67(2):301–20.

44. Torang A, Gupta P, Klinke DJ. An elastic-net logistic regression approach to generate classifiers and gene signatures for types of immune cells and T helper cell subsets. BMC Bioinformatics. 2019 Dec;20(1):433.

45. Cannoodt R, Saelens W, Sichien D, Tavernier S, Janssens S, Guilliams M, et al. SCORPIUS improves trajectory inference and identifies novel modules in dendritic cell development [Internet]. 2016 [cited 2024 Oct 31]. Available from: http://biorxiv.org/lookup/doi/10.1101/079509

46. FlowJo^TM^ Software (for Windows) [Flow Cytometry Analysis]. Ashland, OR: Becton, Dickinson and Company; 2023.

47. RStudio Team. RStudio: Integrated Development Environment for R [Internet]. Boston, MA: RStudio, PBC.; 2020. Available from: http://www.rstudio.com/

48. Robinson JT, Thorvaldsdóttir H, Winckler W, Guttman M, Lander ES, Getz G, et al. Integrative genomics viewer. Nat Biotechnol. 2011 Jan;29(1):24–6.

49. Wickham H. ggplot2: elegant graphics for data analysis. Second edition. Switzerland: Springer; 2016. 260 p. (Use R!).

50. Gu Z. Complex heatmap visualization. iMeta. 2022 Sep;1(3):e43.

51. Samuel Marsh, Maëlle Salmon, Paul Hoffman. samuel-marsh/scCustomize: Version 2.1.2 [Internet]. Zenodo; 2024 [cited 2024 Jul 22]. Available from: https://zenodo.org/doi/10.5281/zenodo.5706430

52. Domínguez J. Workshop studies on monoclonal antibodies in the myeloid panel with CD11 specificity. Veterinary Immunology and Immunopathology. 2001 Jul 20;80(1–2):111–9.

53. Nutt SL, Chopin M. Transcriptional Networks Driving Dendritic Cell Differentiation and Function. Immunity. 2020 Jun;52(6):942–56.

54. Vu Manh TP, Elhmouzi-Younes J, Urien C, Ruscanu S, Jouneau L, Bourge M, et al. Defining Mononuclear Phagocyte Subset Homology Across Several Distant Warm-Blooded Vertebrates Through Comparative Transcriptomics. Front Immunol [Internet]. 2015 Jun 19 [cited 2024 Aug 8];6. Available from: http://journal.frontiersin.org/Article/10.3389/fimmu.2015.00299/abstract

55. Leylek R, Alcántara-Hernández M, Granja JM, Chavez M, Perez K, Diaz OR, et al. Chromatin Landscape Underpinning Human Dendritic Cell Heterogeneity. Cell Reports. 2020 Sep;32(12):108180.

56. Valente M, Collinet N, Vu Manh TP, Popoff D, Rahmani K, Naciri K, et al. Novel mouse models based on intersectional genetics to identify and characterize plasmacytoid dendritic cells. Nat Immunol. 2023 Apr;24(4):714–28.

57. Grajkowska LT, Ceribelli M, Lau CM, Warren ME, Tiniakou I, Nakandakari Higa S, et al. Isoform-Specific Expression and Feedback Regulation of E Protein TCF4 Control Dendritic Cell Lineage Specification. Immunity. 2017 Jan;46(1):65–77.

58. Doulatov S, Notta F, Rice KL, Howell L, Zelent A, Licht JD, et al. PLZF is a regulator of homeostatic and cytokine-induced myeloid development. Genes Dev. 2009 Sep 1;23(17):2076–87.

59. Geijtenbeek TBH, Gringhuis SI. Signalling through C-type lectin receptors: shaping immune responses. Nat Rev Immunol. 2009 Jul;9(7):465–79.

60. Scur M, Parsons BD, Dey S, Makrigiannis AP. The diverse roles of C-type lectin-like receptors in immunity. Front Immunol. 2023 Feb 27;14:1126043.

61. Drouin M, Saenz J, Chiffoleau E. C-Type Lectin-Like Receptors: Head or Tail in Cell Death Immunity. Front Immunol. 2020 Feb 18;11:251.

62. Lee HK, Mattei LM, Steinberg BE, Alberts P, Lee YH, Chervonsky A, et al. In Vivo Requirement for Atg5 in Antigen Presentation by Dendritic Cells. Immunity. 2010 Feb;32(2):227–39.

63. Lei L, Bandola-Simon J, Roche PA. Ubiquitin-conjugating enzyme E2 D1 (Ube2D1) mediates lysine-independent ubiquitination of the E3 ubiquitin ligase March-I. Journal of Biological Chemistry. 2018 Mar;293(11):3904–12.

64. Jancic C, Savina A, Wasmeier C, Tolmachova T, El-Benna J, Dang PMC, et al. Rab27a regulates phagosomal pH and NADPH oxidase recruitment to dendritic cell phagosomes. Nat Cell Biol. 2007 Apr;9(4):367–78.

65. Harden JL, Egilmez NK. Indoleamine 2,3-Dioxygenase and Dendritic Cell Tolerogenicity. Immunological Investigations. 2012 Aug;41(6–7):738–64.

66. Romagnani S. IL4I1: Key immunoregulator at a crossroads of divergent T-cell functions. Eur J Immunol. 2016 Oct;46(10):2302–5.

67. Gliddon DR, Howard CJ. CD26 is expressed on a restricted subpopulation of dendritic cells in vivo. Eur J Immunol. 2002 May;32(5):1472.

68. Zhou X, Khan S, Huang D, Li L. V-Set and immunoglobulin domain containing (VSIG) proteins as emerging immune checkpoint targets for cancer immunotherapy. Front Immunol. 2022 Sep 15;13:938470.

69. Kotwica-Mojzych K, Jodłowska-Jędrych B, Mojzych M. CD200:CD200R Interactions and Their Importance in Immunoregulation. IJMS. 2021 Feb 5;22(4):1602.

70. Reeh H, Rudolph N, Billing U, Christen H, Streif S, Bullinger E, et al. Response to IL-6 trans- and IL-6 classic signalling is determined by the ratio of the IL-6 receptor α to gp130 expression: fusing experimental insights and dynamic modelling. Cell Commun Signal. 2019 Dec;17(1):46.

71. Bachem A, Güttler S, Hartung E, Ebstein F, Schaefer M, Tannert A, et al. Superior antigen cross-presentation and XCR1 expression define human CD11c+CD141+ cells as homologues of mouse CD8+ dendritic cells. Journal of Experimental Medicine. 2010 Jun 7;207(6):1273–81.

72. Dorner BG, Dorner MB, Zhou X, Opitz C, Mora A, Güttler S, et al. Selective Expression of the Chemokine Receptor XCR1 on Cross-presenting Dendritic Cells Determines Cooperation with CD8+ T Cells. Immunity. 2009 Nov;31(5):823–33.

73. Hong W, Yang B, He Q, Wang J, Weng Q. New Insights of CCR7 Signaling in Dendritic Cell Migration and Inflammatory Diseases. Front Pharmacol. 2022 Feb 25;13:841687.

74. Cordero H. Chemokine receptors in primary and secondary lymphoid tissues. In: International Review of Cell and Molecular Biology [Internet]. Elsevier; 2024 [cited 2025 Feb 25]. p. 1–19. Available from: https://linkinghub.elsevier.com/retrieve/pii/S1937644823001673

75. Cytlak U, Resteu A, Pagan S, Green K, Milne P, Maisuria S, et al. Differential IRF8 Transcription Factor Requirement Defines Two Pathways of Dendritic Cell Development in Humans. Immunity. 2020 Aug;53(2):353–370.e8.

76. Sasaki I, Hoshino K, Sugiyama T, Yamazaki C, Yano T, Iizuka A, et al. Spi-B is critical for plasmacytoid dendritic cell function and development. Blood. 2012 Dec 6;120(24):4733–43.

77. Alcántara-Hernández M, Leylek R, Wagar LE, Engleman EG, Keler T, Marinkovich MP, et al. High-Dimensional Phenotypic Mapping of Human Dendritic Cells Reveals Interindividual Variation and Tissue Specialization. Immunity. 2017 Dec;47(6):1037–1050.e6.

78. See P, Dutertre CA, Chen J, Günther P, McGovern N, Irac SE, et al. Mapping the human DC lineage through the integration of high-dimensional techniques. Science. 2017 Jun 9;356(6342):eaag3009.

79. Luber CA, Cox J, Lauterbach H, Fancke B, Selbach M, Tschopp J, et al. Quantitative Proteomics Reveals Subset-Specific Viral Recognition in Dendritic Cells. Immunity. 2010 Feb;32(2):279–89.

80. Yamazaki C, Miyamoto R, Hoshino K, Fukuda Y, Sasaki I, Saito M, et al. Conservation of a chemokine system, XCR1 and its ligand, XCL1, between human and mice. Biochemical and Biophysical Research Communications. 2010 Jul;397(4):756–61.

81. Talker SC, Barut GT, Lischer HEL, Rufener R, Von Münchow L, Bruggmann R, et al. Monocyte biology conserved across species: Functional insights from cattle. Front Immunol. 2022 Jul 29;13:889175.

82. Busch R, Neese RA, Awada M, Hayes GM, Hellerstein MK. Measurement of cell proliferation by heavy water labeling. Nat Protoc. 2007 Dec;2(12):3045–57.

83. Patel AA, Zhang Y, Fullerton JN, Boelen L, Rongvaux A, Maini AA, et al. The fate and lifespan of human monocyte subsets in steady state and systemic inflammation. Journal of Experimental Medicine. 2017 Jul 3;214(7):1913–23.

84. Esashi E, Wang YH, Perng O, Qin XF, Liu YJ, Watowich SS. The Signal Transducer STAT5 Inhibits Plasmacytoid Dendritic Cell Development by Suppressing Transcription Factor IRF8. Immunity. 2008 Apr;28(4):509–20.

85. Edwards JC, Everett HE, Pedrera M, Mokhtar H, Marchi E, Soldevila F, et al. CD1− and CD1+ porcine blood dendritic cells are enriched for the orthologues of the two major mammalian conventional subsets. Scientific Reports. 2017 Jan 20;7(1):40942.

86. Tussiwand R, Rodrigues PF. Where’s Waldo: Identifying DCs within Mononuclear Phagocytes during Inflammation. Immunity. 2020 Jun;52(6):892–4.

87. Barut GT, Kreuzer M, Bruggmann R, Summerfield A, Talker SC. Single-cell transcriptomics reveals striking heterogeneity and functional organization of dendritic and monocytic cells in the bovine mesenteric lymph node. Front Immunol. 2023 Jan 6;13:1099357.

