## Supplementary Figures for "Unraveling dendritic-cell diversity in blood of pigs: welcome tDC and DC3"

**Supp. Fig. 1:** Complete gating strategy for the phenotypic characterization of the unknown DC subset in porcine blood.

**Supp.Fig. 2:** Bulk RNA-seq of sorted DC subsets.

**Supp. Fig. 3:** Bulk transcriptomic signatures of DC subsets, including pptDC, related to immune functions.

**Supp.Fig. 4:** Clusters (scRNA-seq) excluded from further analyses based on quality control and B-cell-/ NK-cell-annotated signatures.

**Supp. Fig. 5:** Complete gating strategy related to Figure 4E.

**Supp. Fig. 6:** Gene set enrichment analysis of bulk-RNA-seq-derived gene sets in scRNA-seq clusters.

**Supp. Fig. 7:** Analysis of a published scRNA-seq dataset of human blood DC.

**Supp. Fig. 8:** Trajectory analysis for cells bridging tDC and cDC2 in scRNA-seq clustering.

### Supplementary Figure 1

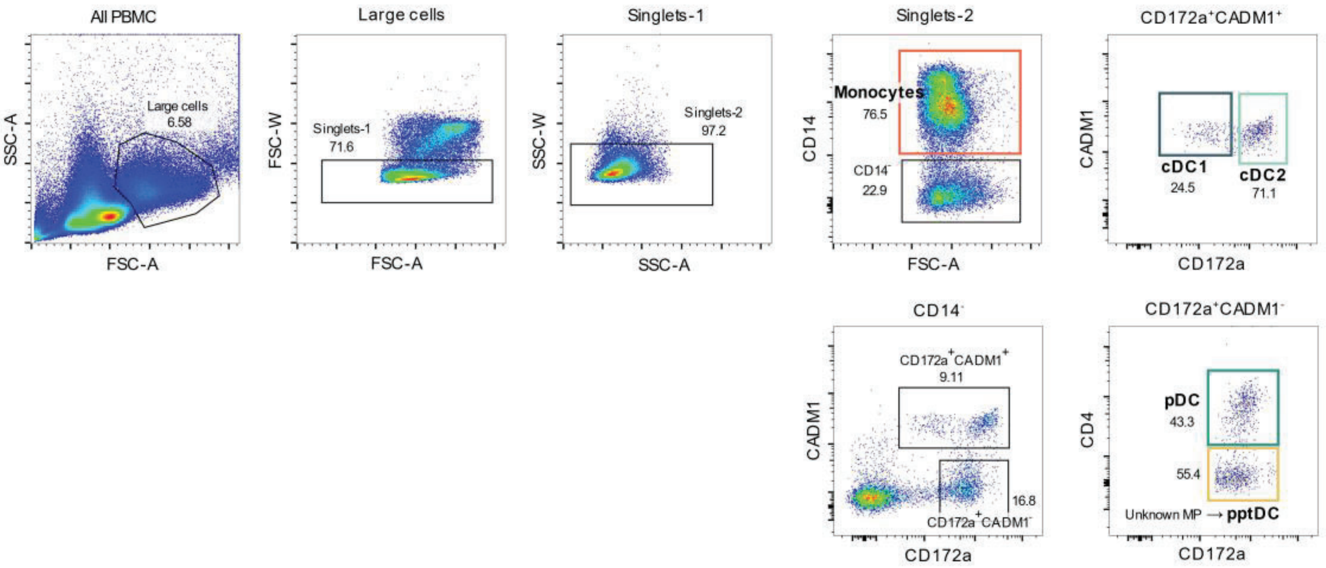


**Supplementary Figure 1: Complete gating strategy for the phenotypic characterization of the unknown DC subset in porcine blood.** PBMC were isolated from the blood of four 12 to 24-month-old pigs and stained for flow cytometry. Representative gating for mononuclear phagocyte (MP) subsets. Following selection of large cells ($SSC^{high}FSC^{high}$) and doublet exclusion based on forward scatter (FSC-A vs. FSC-W) and side scatter (SSC-A vs. SSC-W) characteristics, $CD14^{+}$ cells were defined as monocytes and four subsets were distinguished among $CD14^{-}$ cells based on the surface expression of CADM1, CD172a and CD4: cDC1 identified as $CD172a^{low}CADM1^{+}$ cells, cDC2 as $CD172a^{+}CADM1^{+}$ cells, pDC as $CD172a^{+}CADM1^{-}CD4^{+}$ cells and a newly described DC subset as $CD172a^{+}CADM1^{-}CD4^{-}$ cells.

### Supplementary Figure 2

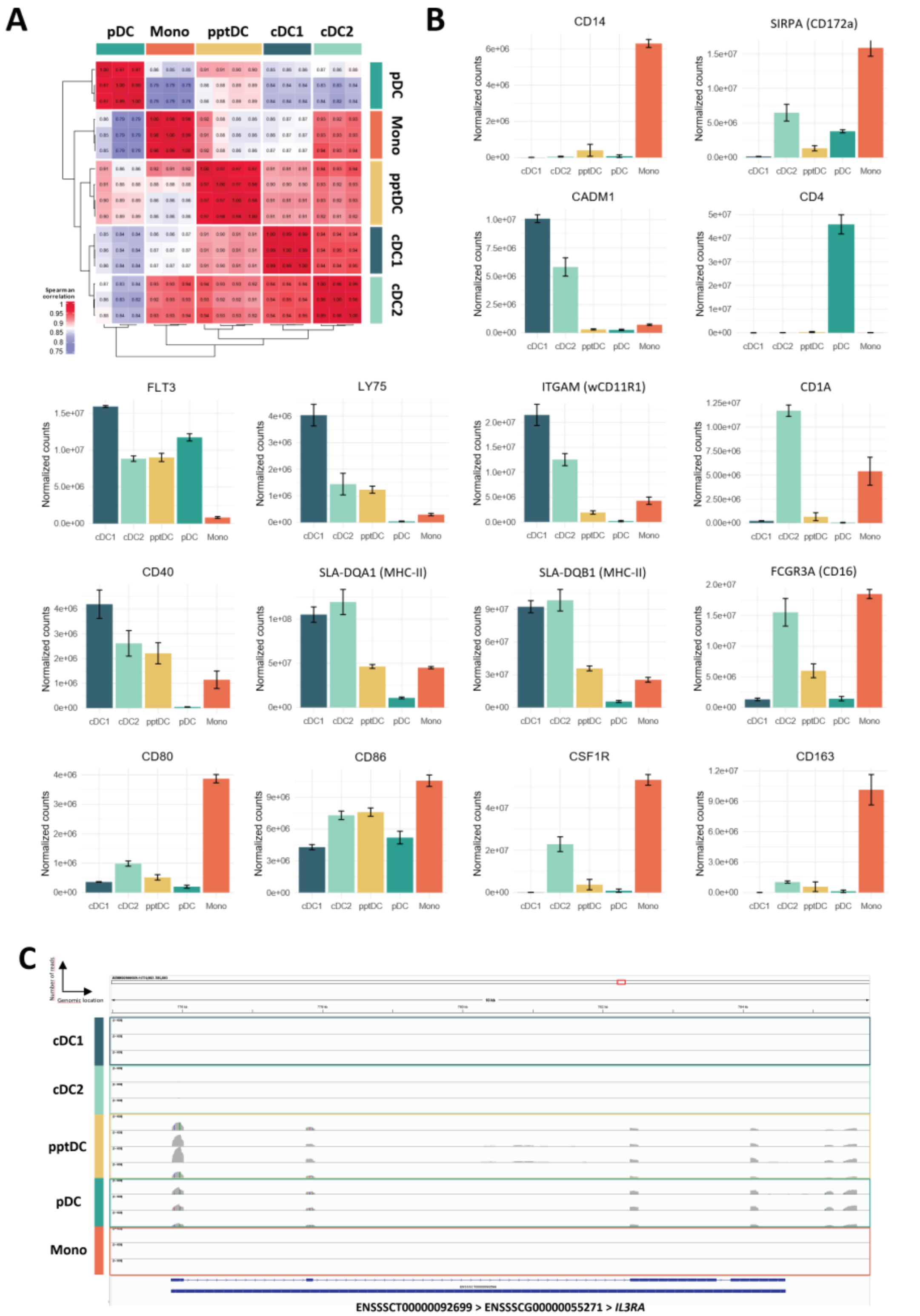


**Supplementary Figure 2: Bulk RNA-seq of sorted DC subsets.** (**A**) MP subset-to-subset correlation analysis based on normalized gene expression data using the Spearman correlation method, with hierarchical clustering based on Spearman distances. (**B**) Transcription of genes encoding sixteen cell-surface markers analyzed by flow cytometry. Bar plots show normalized counts for each gene and MP subset (mean ± standard deviation). (**C**) Read coverage for IL3RA (ENSSSCG00000055271), evidencing expression of this gene in pptDC and pDC. “Mono” refers to “monocytes”.

### Supplementary Figure 3

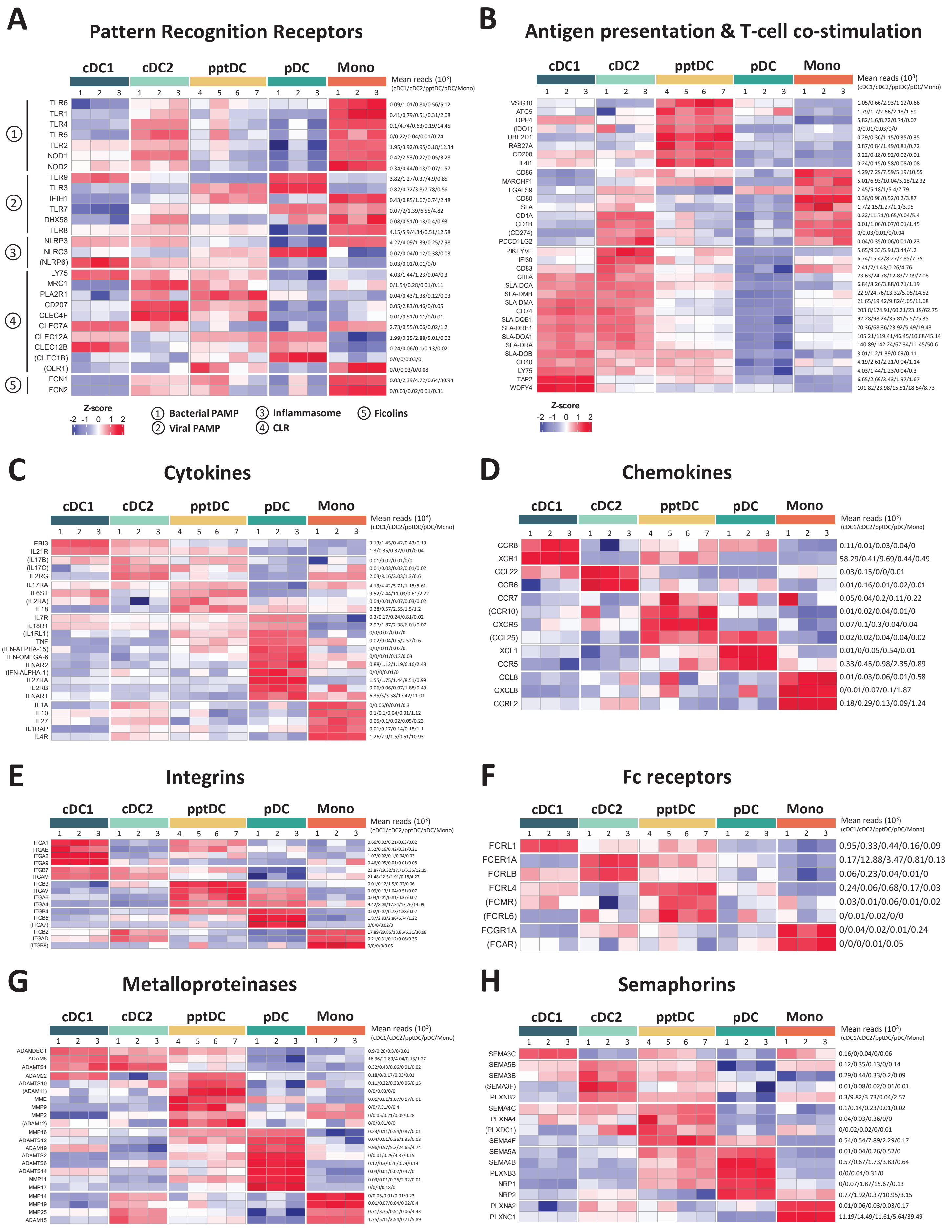

**Supplementary Figure 3: Bulk transcriptomic signatures of DC subsets, including pptDC, related to immune functions.** Bulk RNA-seq was performed on five sorted mononuclear phagocyte (MP) subsets. Heatmaps show gene expression for selected categories. Mean kilo reads are given to the right of each heatmap. Genes in parenthesis indicate a mean of < 100 reads across all subsets and should be interpreted with caution. “Mono” refers to “monocytes”.

### Supplementary Figure 4

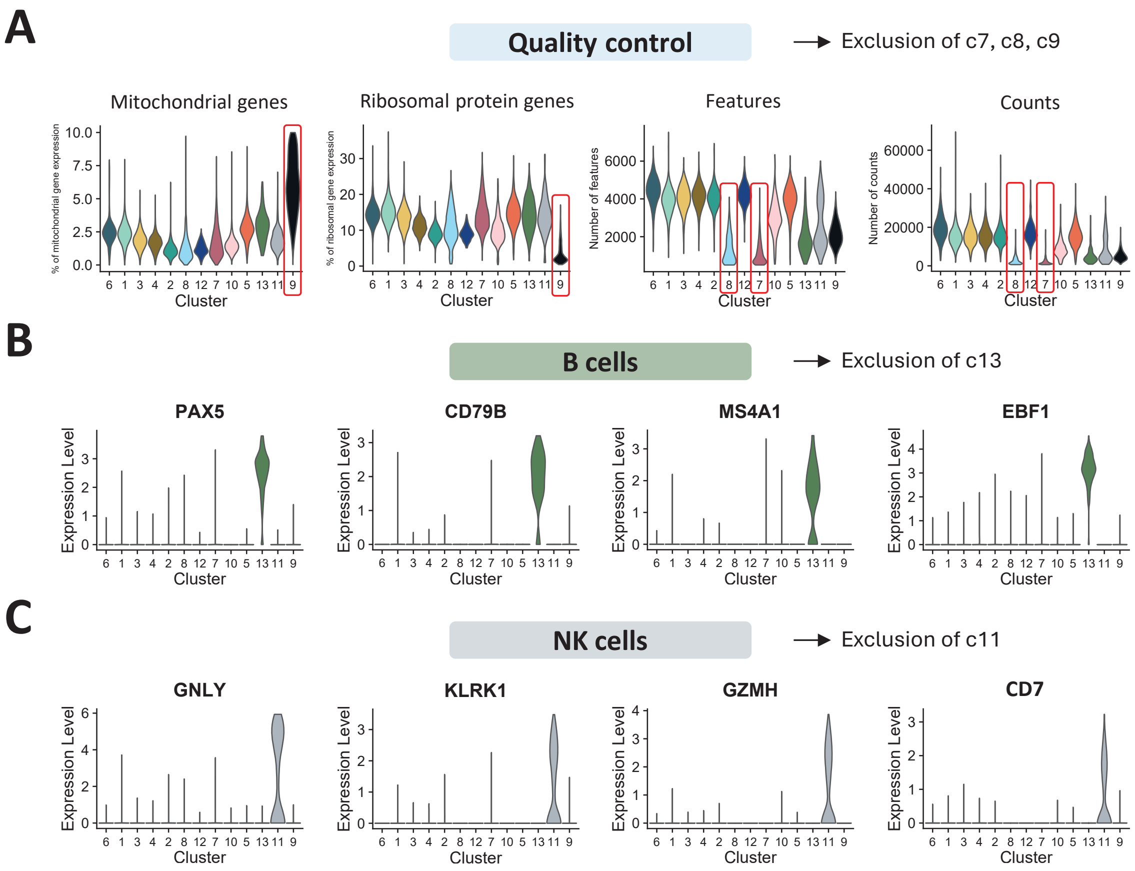


**Supplementary Figure 4: Clusters (scRNA-seq) excluded from further analyses based on quality control and B-cell-/ NK-cell-annotated signatures.** Dendritic cells (DC) were sorted from PBMC of three pigs and subjected to 10x Genomics scRNA-seq. Data from approximately 10 000 DC per sample was analyzed. (**A**) Quality control . Red frames indicate excluded clusters (c7, c8, c9). (**B**) B-cell associated gene expression detected in excluded c13. (**C**) NK-cell associated gene expression detected in excluded c11.

### Supplementary Figure 5

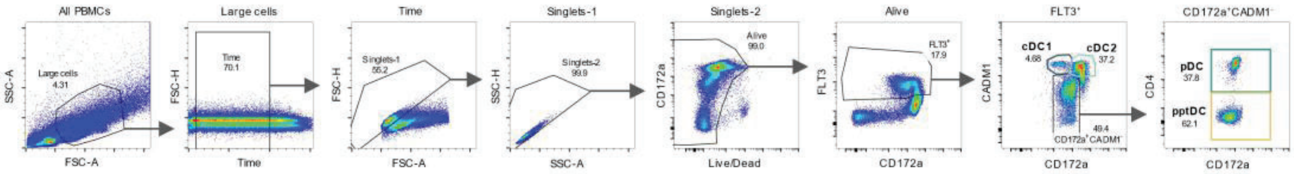


**Supplementary Figure 5: Complete gating strategy related to Figure 4E.** Freshly isolated PBMC of the three animals included in the scRNA-seq analysis were stained for flow cytometry. Following selection of large cells ($SSC^{high}FSC^{high}$), time-gating, doublet exclusion based on forward scatter (FSC-A vs. FSC-H) and side scatter (SSC-A vs. SSC-H) characteristics, and exclusion of dead cells ($Live/Dead^{-}$), $Flt3^{+}$ cells were defined as DC. The four DC subsets were distinguished based on the surface expression of CADM1, CD172a and CD4: cDC1 identified as $CD172a^{low}CADM1^{+}$ cells, cDC2 as $CD172a^{+}CADM1^{+}$ cells, pDC as $CD172a^{+}CADM1^{-}CD4^{+}$ cells and pptDC as $CD172a^{+}CADM1^{-}CD4^{-}$ cells.

### Supplementary Figure 6

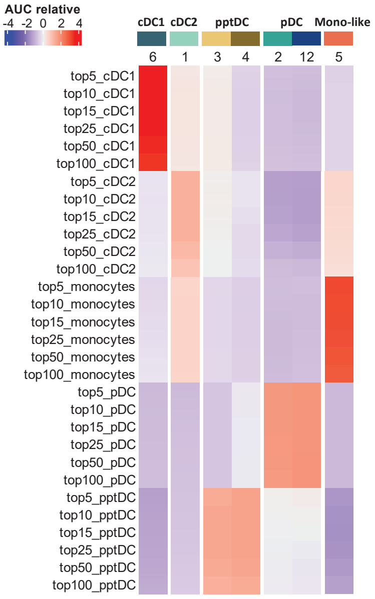


**Supplementary Figure 6: Gene set enrichment analysis of bulk-RNA-seq-derived gene sets in scRNA-seq clusters.** Sorting of porcine mononuclear phagocyte (MP) subsets was performed as shown in **Figure 2A.** Gene sets were obtained from DESeq2 analysis of bulk-sequenced subsets. Gene set enrichment scores were calculated with the AUCell method based on the top 5, 10, 15, 25, 50 and 100%expressed genes in each single cell, and the gene sets. Heatmap shows averaged scaled AUC scores calculated for each cluster. Higher AUC scores correspond to a higher percentage of genes from a gene set being detected in the top expressed genes in a cell, defined at different levels. "Mono-like" refers to "monocyte-like cells".

### Supplementary Figure 7

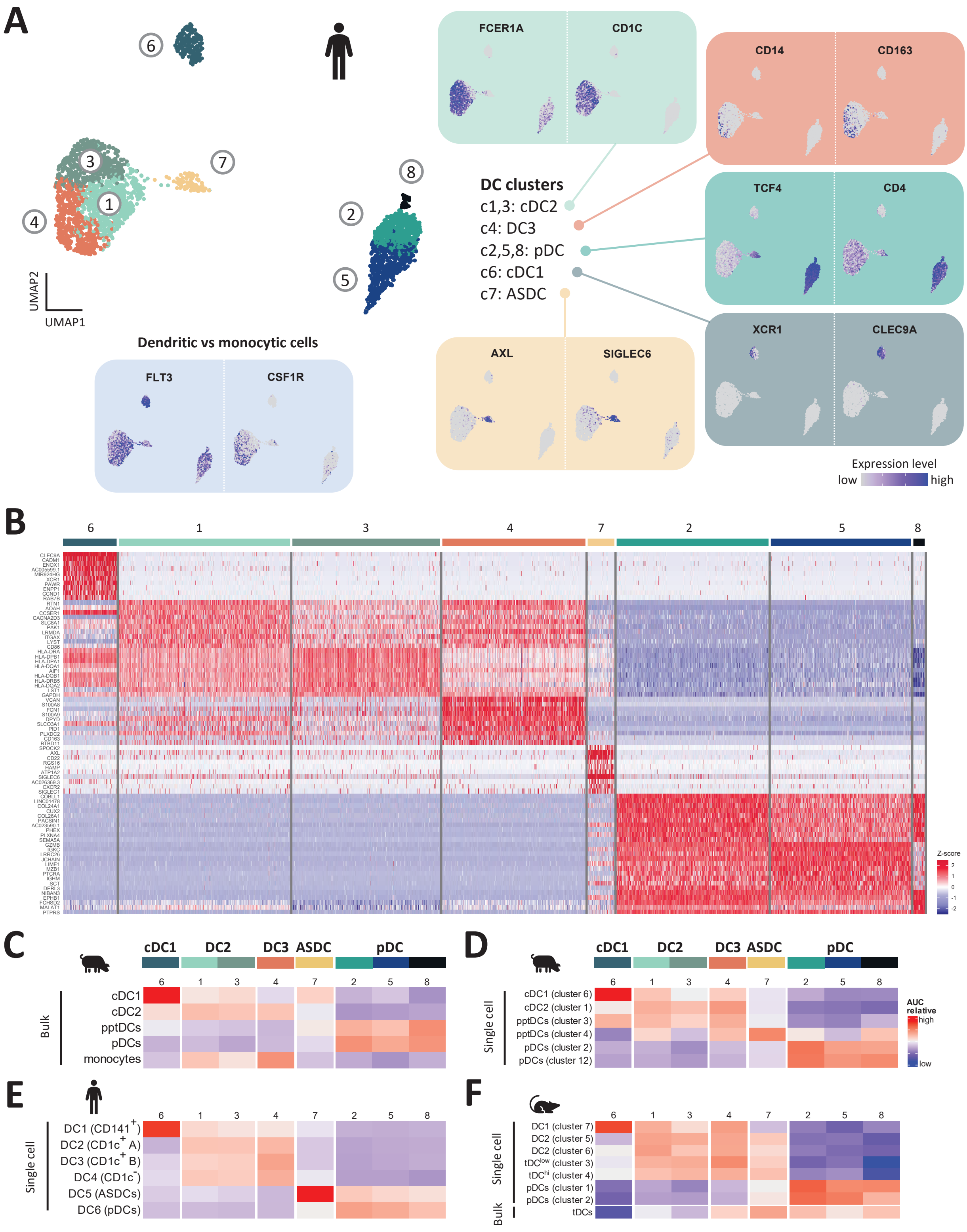


**Supplementary Figure 7: Analysis of a published scRNA-seq dataset of human blood DC.** Analysis of the scRNA-seq dataset of human blood dendritic cells (DC) generated by Lubin *et al.*[39] (approximately 3,000 cells). DC were sorted by FACS and subjected to 10x Genomics scRNA-seq. (**A**) Data were processed (see **Material and Methods** section) and a clustering was performed with a resolution of 0.8 (Leiden algorithm), resulting in 8 different clusters visualized by UMAP plot. Feature plots showing the expression of *FLT3* and *CSF1R*, distinguishing between dendritic (clusters 1, 2, 3, 5, 6, 7 and 8) and monocytic cell clusters (cluster 4), respectively, and the expression of key genes specific for cDC2 (*FCER1A* and *CD1C*), cDC1 (*XCR1* and *CLEC9A*), pDC (*CD4* and *TCF4*), DC3 (*CD163* and *CD14*) and ASDC (*AXL* and *SIGLEC6*). (**B**) Heatmap showing the top 10 differentially expressed genes (p_val_adj) in each cluster, as determined by Seurat's *FindAllMarkers()* function. Complete gene lists are available as **Supp. Data 3**.

### Supplementary Figure 8

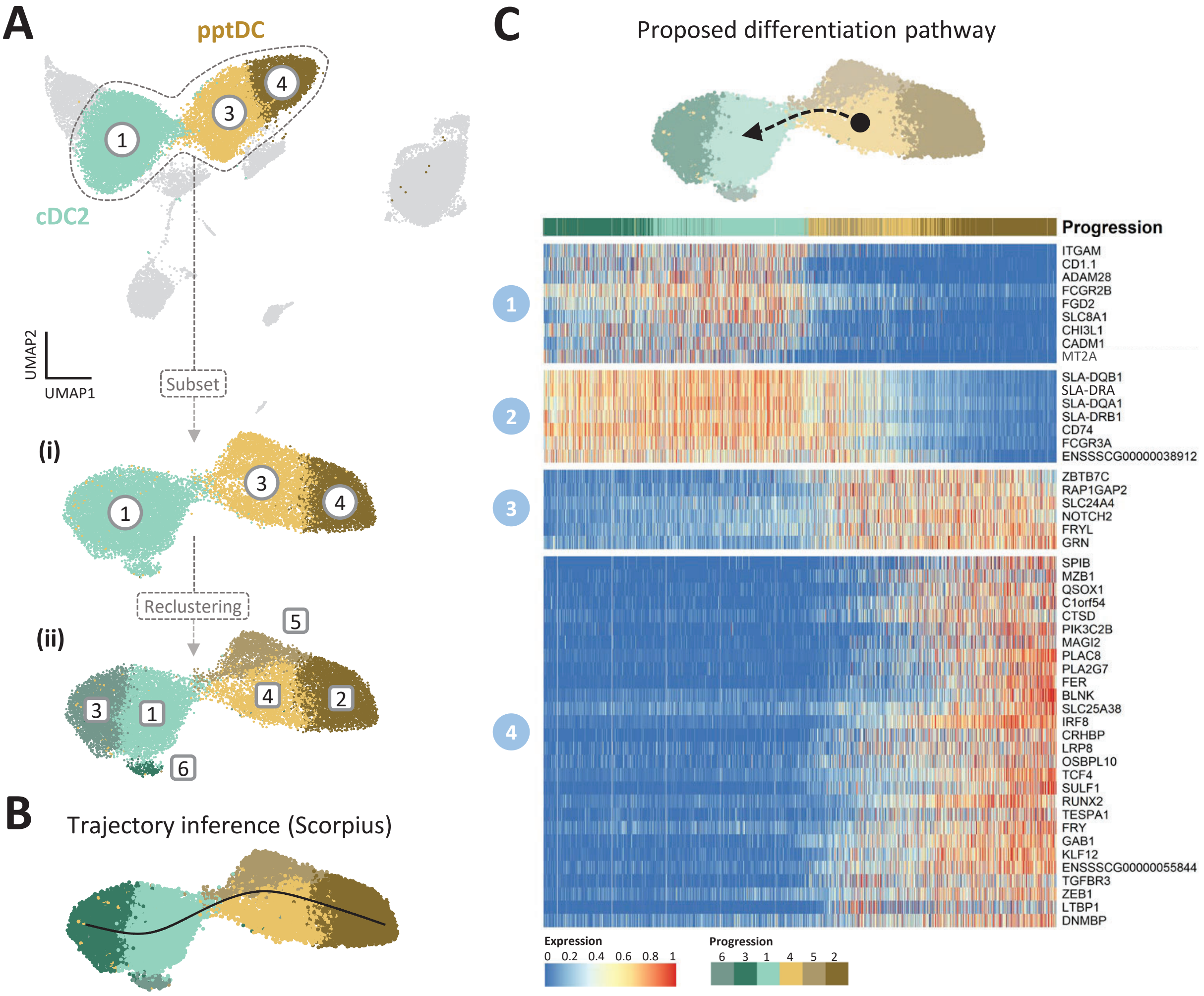


**Supplementary Figure 8: Trajectory analysis for cells bridging tDC and cDC2 in scRNA-seq clustering.** (**A**) A subset of cells belonging to the clusters 1, 3 and 4 (approximately 17 400 cells) was generated from the initial scRNA-seq dataset (see **Material and Methods** section for details) (**i**) and a new clustering was performed with a resolution of 0.6 (Leiden algorithm), resulting in 6 distinct clusters visualized by UMAP plot (**ii**). Differentially expressed genes as determined by Seurat's *FindAllMarkers()* function are included in **Supp. Data 3**. (**B**) A trajectory inference analysis was performed with Scorpius, and cells were ordered along the inferred linear trajectory visualized by UMAP plot. (**C**) Heatmap showing normalized expression values (scaled from 0 to 1) of the top 50 important genes along the inferred trajectory. Genes were grouped into four modules (1-4) by Scorpius, based on their distinct patterns of up- and down-regulation across the different trajectory waves. Complete gene list is available as **Supp. Data 3**. On top, proposed differentiation pathway illustrating the transition from cluster 3 to cluster 1.
